# Multi-omics Reveals Immune Response and Metabolic Profiles during High-Altitude Mountaineering

**DOI:** 10.1101/2024.05.03.592361

**Authors:** Jianhua Yin, Jingzhi Lv, Shichen Yang, Yang Wang, Zhuoli Huang, Xue Wang, Guixue Hou, Wenwen Zhou, Ying Liu, Weikai Wang, Xiumei Lin, Yunting Huang, Yuhui Zheng, Chen Wei, Yue Yuan, Yaling Huang, Chang Liu, Haoran Tao, Huanhuan Liu, Ruquan Liu, Yan Zhang, Guodan Zeng, Peng Gao, Longqi Liu, Jun Cao, Chuanyu Liu, Xin Jin, Jian Wang

## Abstract

The physiological perturbations induced by high-altitude exposure in mountain climbers, manifesting as metabolic and immunologic deviations, have been previously reported but are not fully understood. In this study, we obtained longitudinal multi-omic profiles of blood samples for healthy mountain climbers during two mountaineering stages (acclimatization and extreme altitude mountaineering). Our integrative assay included metabolomics and lipidomics profiling of plasma coupled with single-cell transcriptomic analysis of 375,722 immune cells. Longitudinal analysis revealed dynamic immune response profiles, during the acclimatization period, characterized by the downregulation of inflammatory responses in monocytes and classical dendritic cells (cDCs) and an increase in the proportion of cytotoxic CD8^+^ T cells with enhanced immune effector processes. In contrast, during extreme altitude mountaineering, the activation of inflammatory responses and impairment of T cell effector function were observed, concomitant with an increased cellular response to hypoxia and oxidative stress pathways. Furthermore, we found upregulated glycolysis and antioxidant gene expression during extreme altitude mountaineering, which was primarily orchestrated by *HIF1A* and *NFE2L2*, while decreased expression of these genes was observed in dysregulated plasmacytoid dendritic cells (pDCs). Finally, high-resolution plasma metabolic analysis revealed significant alterations in the metabolism of climbers, involving enhanced glutamine and fatty acid metabolism.

## INTRODUCTION

High-altitude mountaineering, with the main objective of reaching a summit, is a challenging and potentially dangerous activity. Increased accessibility to high mountains has led to the growing popularity of mountaineering[1]. Mountain climbers are confronted with harsh environmental conditions upon exposure to high altitudes, among which the most crucial variable is hypoxia owing to the progressive decline in barometric pressure with increasing altitude[2]. Several physiological responses are initiated in climbers to adapt to reduced oxygen availability, such as enhanced ventilatory response, increased cardiac output, and improved tissue oxygen extraction[3]. The combination of hypoxia and intense physical activity may impose greater physiological and metabolic demands on mountain climbers[4]. Notably, mountain climbers are susceptible to a range of health issues, including altitude illness[3], body mass loss[5], immune dysfunction and increased vulnerability to infections[6]. Among them, immunological and metabolic outcomes are key mediators, but the underlying mechanisms remain incompletely characterized.

Moderate exercise is beneficial for immune function, while an excessive duration and intensity of exercise may lead to immunological impairment[7, 8]; therefore, it is crucial to understand the immune outcomes of different types of exercise. Previous studies have reported that low oxygen availability at high altitudes has a significant impact on the abundance and function of immune cells; specifically, it leads to the downregulated expression of innate immune-related genes[9], a reduction in the proportion of plasmacytoid dendritic cells (pDCs)[10], and a decrease in CD3^+^ T lymphocytes[11]. Importantly, exercise at high altitudes may intensify the extent of hypoxia, further impeding immune function[4] and elevating the levels of inflammation and oxidative stress[12–14]. Notably, exercise has the potential to reprogram immune cell metabolism through alterations in oxygen availability, the regulation of cellular energy sensors, and the accumulation of immunometabolic regulators[4, 15]. Previous studies have assessed the metabolic activity of immune cells during high-altitude exercise by assessing metabolism-related parameters, including reactive oxygen species (ROS), mitochondrial activity, and intracellular calcium ions[16, 17]. These metabolic changes may subsequently impact the immune cell state and cell fate[18]. However, the underlying mechanism of the immune response to high-altitude mountaineering remains elusive. In addition, in a previous study, participants who ascended to high altitudes exhibited substantial alterations in plasma metabolic molecules[19]. A comprehensive characterization of both immune cells and plasma analytes from mountain climbers is needed to fully understand the dynamic alterations in the immune response and plasma metabolites.

The emergence of single-cell RNA sequencing (scRNA-seq) has facilitated an in-depth dissection of immune response changes driven by exercise and has been applied to exercise immunology[20]. To elucidate the immune response and metabolic alterations during high-altitude mountaineering, we performed multi-omics profiling, including targeted metabolomics and lipidomics of plasma, as well as single-cell transcriptome analysis of circulating immune cells, on each sample from eleven mountain climbers (Figure 1A). In this study, our longitudinal analysis revealed dynamic immune responses at different stages of high-altitude mountaineering. In particular, we identified a crucial immune adaptive response mediated by the transcription factors (TFs) *HIF1A* and *NFE2L2* during extreme altitude mountaineering. Furthermore, substantial alterations in plasma metabolites, including increased glutamine metabolism and fatty acid metabolism, were also observed during high-altitude mountaineering. Overall, the results from this study improve our understanding of the dynamic immune and metabolic landscapes elicited by high-altitude mountaineering.

**Figure 1.**
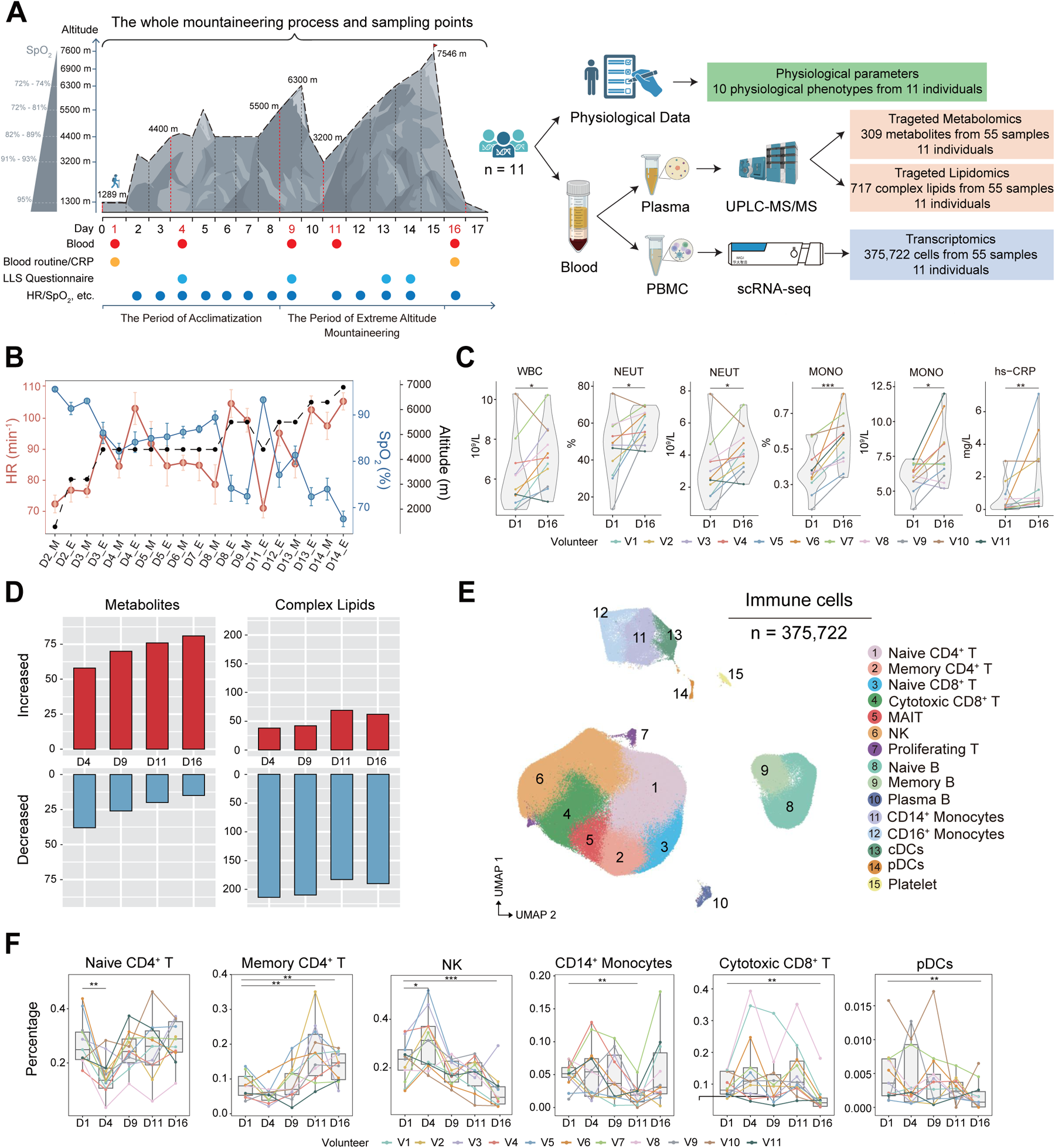
Overview of the multi-omics profiling of metabolic molecules and immune responses in mountain climbers. (A) Flowchart depicting the overall mountaineering design, sample collection and multi-omics data analysis. All the data were collected from 11 healthy mountain climbers. The figures were created with Adobe Illustrator 2023 and BioRender.com. (B) Effects of high altitude on heart rate (HR) and blood saturation (SpO_2_) during mountaineering. Red represents HR, blue represents SpO_2_, and the black dotted line represents altitude. M was measured in the morning, and E was measured in the evening. (C) Violin plots showing significant changes in selected routine blood parameters and C-reactive protein (CRP) levels before (D1) and after (D16) the mountaineering expedition. The two-sided *p* values from the paired Wilcoxon rank-sum test are shown, \**p* < 0.05, \*\**p* < 0.01, \*\*\**p* < 0.001. (D) Bar plots showing changes in the numbers of significantly differentially abundant metabolites and complex lipids during mountaineering. (E) Uniform manifold approximation and projection (UMAP) plot showing 15 cell types among 375,722 circulating immune cells. Cells are colored according to different cell subtypes. (F) Boxplots showing the relative proportion changes in selected subsets during mountaineering (day 1, day 4, day 7, day 11 and day 16, n = 11); sample points from the same mountain climber are connected with the same color. The two-sided *p* values from the paired Wilcoxon rank-sum test are shown, \**p* < 0.05, \*\**p* < 0.01, \*\*\**p* < 0.001.

## RESULTS

### Effects of high altitude on physiological parameters in mountain climbers

This study was conducted during an expedition to the Muztagh Ata peak (7546 m) with the aim of investigating the dynamic physiological, metabolic, and immune responses of mountain climbers at high altitudes. Eleven climbers (8 males, 3 females) with a mean age of 37.4 ± 11.4 years and a mean body mass index (BMI) of 23.60 ± 3.16 kg/m^2^ were recruited (Table S1). After reaching 1289 m on D1, the mountaineering experimental plan consisted of two stages: the period of acclimatization (D2-D8) and then the period of extreme altitude mountaineering (D9-D16) (Table S1). A total of 10 physiological phenotypic indexes, including blood oxygen saturation (SpO_2_), heart rate (HR), Lake Louise score (LLS), body composition, and routine blood and ultrasensitive C-reactive protein (CRP) tests, were assessed for eleven climbers (Figure 1A). Furthermore, 55 peripheral blood samples were collected longitudinally at five different time points of mountaineering: D1 (altitude, 1289 m; SpO_2_, 95%), performed in low altitudes as a baseline control; D4 (altitude, 4400 m; SpO_2_, 82%) and D9 (altitude, 5500 m; SpO_2_, 72%), performed following acclimatization to high altitudes of 4400 m and 5500 m; and D11 (altitude, 3200 m) and D16 (altitude, 1289 m), performed after reaching extreme altitudes of 6300 m and 7546 m (Table S1).

We first analyzed the variations in SpO_2_ and HR during different stages of high-altitude mountaineering. Overall, we observed a progressive decrease in the SpO_2_ of climbers with increasing altitude and a compensatory increase in the HR from 72 to 105 min^-1^, with high consistency among climbers (Figure 1B and S1A). When the mountain climbers were exposed to extreme altitudes of 6,300 m and 6,900 m, the SpO_2_ decreased to 72% and 69%, respectively. However, in comparison to the initial exposure at 4400 m on D3, we observed a significant increase in SpO_2_ from 86% to 93% on D11 following acclimatization, accompanied by a notable decrease in HR from 94 min^-1^ to 71 min^-1^; such trends were also observed upon re-exposure to an altitude of 5,500 m (Figure S1B). These data indicated that the mountain climbers may have acclimatized to the reduced oxygen levels at very high altitudes (3,500-5,500 m) following the acclimatization period (D2-D8). Next, we compared the changes in body composition characteristics, including BMI, body fat percentage (BFP%), muscle mass (MM, kg) and fat mass (FM, kg), and no significant changes were observed in these parameters (Figure S1C). In addition, routine blood tests and CRP analysis were performed before (D1) and after (D16) the mountaineering expedition (Figure 1A). The results showed a significant increase in the numbers of neutrophils (NEUT) and monocytes (MONO) on D16 compared with those at baseline (D1) (Figure 1C). Moreover, we detected a significant increase in the level of CRP (Figure 1C), a marker of the inflammatory response, consistent with previous reports that long and arduous trekking at high altitudes leads to an increase in the inflammatory response[5]. Notably, the proportion of lymphocytes significantly decreased on D16 (Figure S2). In general, the mountaineering expedition induced dynamic changes in multiple physiological phenotypes, which initially confirmed the hypoxic state of mountain climbers at high altitudes and further reflected the impact of high-altitude mountaineering on immune responses.

### Multi-omics profiling of plasma and peripheral immune cells in mountain climbers

Whole blood samples were collected from eleven healthy volunteers at five different time points during mountaineering. Subsequently, the blood samples were separated: the upper plasma was subjected to metabolomic and lipidomic analyses, and the isolated immune cells were subjected to scRNA-seq analysis (Figure 1A). We analyzed the plasma metabolome and lipidome to obtain a global view of the body’s metabolic status, and targeted analysis was performed via ultra-performance liquid chromatography-tandem mass spectrometry (UPLC-MS/MS). After data preprocessing, a total of 309 metabolites and 717 complex lipids were measured per sample, and the list containing all the metabolic analytes can be found in Table S2 and S3. Then, our focus shifted toward metabolites and lipids that were significantly affected by high-altitude mountaineering. The variable importance in projection (VIP) score was used to estimate the differentially abundant metabolites and lipids with the greatest variation in the data; ultimately, 348 significantly changed analytes were selected. Extensive perturbations of these molecules were observed at different time points (Figure 1D). Overall, the preliminary findings indicated that the metabolism of climbers underwent substantial alterations.

We next performed a single-cell transcriptomics analysis of circulating immune cells obtained from mountain climbers to comprehensively characterize the alterations in the immune response induced by mountaineering. After quality control, a total of 375,722 single cells were obtained with an average of 1,083 genes per cell (Figure S3A). The number of cells in each sample ranged from 48,686 to 100,271, and the number of cells at each time point ranged from 12,853 to 55,231 (Figure S3B). Then, uniform manifold approximation and projection (UMAP) for dimensionality reduction was employed, and we manually annotated cells based on the expression of classical markers (Figure S4A). In total, 15 major cell types were identified, namely, naive CD4^+^ T cells, memory CD4^+^ T cells, naive CD8^+^ T cells, cytotoxic CD8^+^ T cells, mucosal-associated invariant T (MAIT) cells, natural killer (NK) cells, proliferating T cells, naive B cells, memory B cells, plasma B cells, CD14^+^ monocytes, CD16^+^ monocytes, classical dendritic cells (cDCs), pDCs and platelets (Figure 1E). Next, we calculated changes in the relative proportions of these major cell subpopulations among the circulating immune cells at each time point. The proportion of naive CD4^+^ T cells decreased significantly on D4, accompanied by a sustained increase in the proportion of memory CD4^+^ T cells on D11 and D16 (Figure 1F). In contrast, the abundance of NK cells initially exhibited a significant increase on D4, followed by a gradual decrease (Figure 1F). Several subpopulations also exhibited notable reductions at later time points, including the proportion of CD14^+^ monocytes on D11 and the proportions of cytotoxic CD8^+^ T cells and pDCs on D16 (Figure 1F, S4B-S4D). These results indicated that the composition of circulating immune cells in mountain climbers significantly changed during high-altitude mountaineering.

### Altered inflammatory response in monocytes and DCs

Previous studies have demonstrated that high-altitude mountaineering induces an increase in the inflammatory response[12]. Considering the critical roles played by myeloid cells in inflammation, we initially investigated their transcriptional alterations during mountaineering. All monocytes and DCs were subclustered (n=33,577) and classified into nine subpopulations based on the expression of classical marker genes, including three CD14^+^ monocyte (CD14^+^ Mono) subgroups: FCN1^+^ CD14^+^ Monos, CCL3^+^ CD14^+^ Monos and IFN^+^ CD14^+^ Monos; three CD16^+^ monocyte (CD16^+^ Mono) subgroups: IFN^+^ CD16^+^ Monos, CD16^+^ Monos and C1^+^ CD16^+^ Monos; and three DC subgroups: cDC1s, cDC2s and pDCs (Figure 2A and 2B). Remarkable alterations in the relative proportions of these subgroups were observed: CCL3^+^ CD14^+^ Monos significantly decreased on D4 and D9, followed by a subsequent recovery and significant expansion on D16, and FCN1^+^ CD14^+^ Monos significantly decreased on D11 and D16 (Figure 2C). The proportion of CD16^+^ Monos significantly increased on D4 and D9, followed by a decreasing trend. Furthermore, the proportion of cDC2s significantly increased on D11, while the proportion of pDCs significantly decreased on D16 (Figure 2C, S5A and S5B).

**Figure 2.**
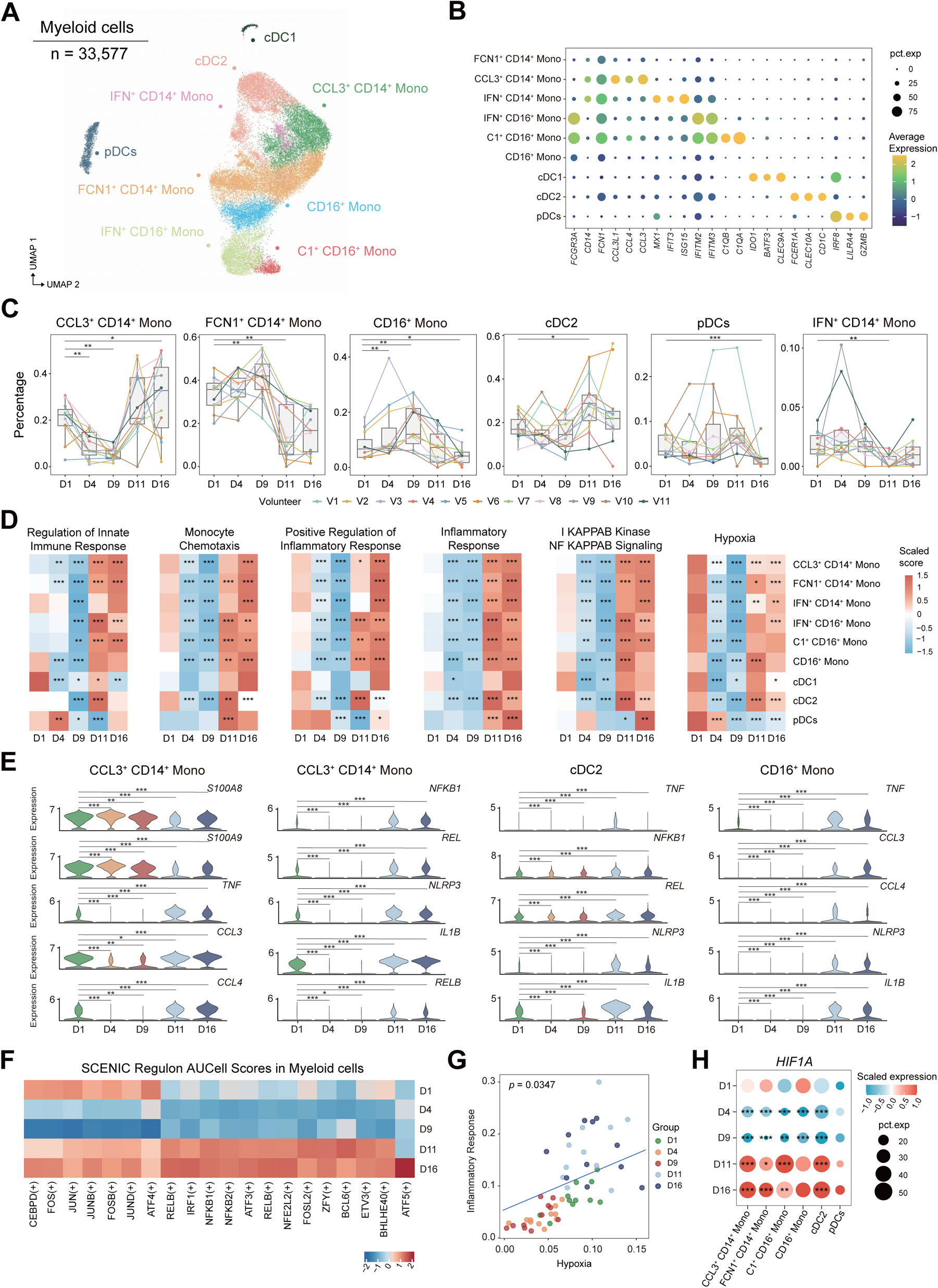
Dynamic changes in the inflammatory response in monocytes and DCs. (A) UMAP plot showing nine myeloid cell subsets. (B) Dot plots depicting the percentages and average expression levels of the canonical genes in nine subsets of myeloid cells. (C) Boxplots showing the changes in the relative proportions of selected myeloid cell subsets before and during mountaineering (day 1, day 4, day 7, day 11 and day 16, n = 11); sample points from the same mountain climber are connected with the same color. The two-sided *p* values from the paired Wilcoxon rank-sum test are shown, \**p* < 0.05, \*\**p* < 0.01, \*\*\**p* < 0.001. (D) Heatmap showing the scaled expression of genes in important biological pathways mediated by myeloid cells across different time points. The two-sided *p* values from the Wilcoxon rank-sum test are shown, \**p* < 0.05, \*\**p* < 0.01, \*\*\**p* < 0.001. (E) Violin plots showing dynamic changes in the expression of selected differentially expressed genes (DEGs) in CCL3^+^ CD14^+^ Monos, cDC2s and CD16^+^ Monos (|log_2_FC| ≥ 0.7, adjusted *p* < 0.05). \**p* < 0.05, \*\**p* < 0.01, \*\*\**p* < 0.001. (F) Heatmap showing the regulon areas under the curve per cell (AUCell) scores for myeloid cells across the five time points, calculated using the SCENIC algorithm. (G) Correlation analysis between hypoxia scores and inflammatory response scores for each climber at all five time points (linear mixed model). (H) Dot plots showing the scaled expression and percentages of *HIF1A* in myeloid cell subsets across different time points. The two-sided *p* values from the Wilcoxon rank-sum test are shown, \**p* < 0.05, \*\**p* < 0.01, \*\*\**p* < 0.001.

To gain insight into the functional changes in different monocyte and DC subgroups during mountaineering, we performed differentially expressed genes (DEGs) analysis (D1 was used as the control group) (Table S4). Gene Ontology (GO) enrichment analysis of these DEGs revealed significant enrichment in pathways related to innate immunity, cytokine production, cellular chemotaxis, hypoxia and the inflammatory response in CCL3^+^ CD14^+^ Monos, FCN1^+^ CD14^+^ Monos, CD16^+^ Monos and cDC2s (Figure S5C). We also performed pathway score analysis to investigate the dynamic changes in key pathways in each myeloid cell subgroup during mountaineering. The expression of genes involved in the innate immune response, inflammatory response, NF-κB signaling and hypoxia pathway was downregulated on D4 and D9, followed by significant upregulation on D11 and D16 in most myeloid cell subgroups, except for pDCs (Figure 2D). The expression of *S100A8* and *S100A9* in CCL3^+^ CD14^+^ Monos was significantly upregulated on D4 and then downregulated (Figure 2E). *S100A8* and *S100A9* have been identified as important endogenous damage-associated molecular pattern (DAMP) molecules, which are early amplifiers of the innate immune response to danger signals[21]. Additionally, the expression of multiple genes involved in inflammation were significantly downregulated on D4 and D9 but upregulated on D11 and D16 in CCL3^+^ CD14^+^ Monos, CD16^+^ Monos, cDC2s and FCN1^+^ CD14^+^ Monos, including cytokine-related genes (*TNF*, *CCL3* and *CCL4*); central transcription factors of inflammatory response NF-κB subunits (*NFκB1* and *REL*); and inflammasome activation-associated genes (*NLRP3* and *IL1B*) (Figure 2E and S5D). The activation of the NLRP3 inflammasome is triggered by a variety of upstream signals, including mitochondrial dysfunction, metabolic changes and ROS[22]. These observations suggested that high-altitude mountaineering induced dynamic changes in inflammatory responses.

Next, we analyzed the dynamic changes in TF regulon activity in monocytes and DCs at different stages using single-cell regulatory network inference and clustering (SCENIC)[23]. The regulon activities of multiple TFs, such as inflammation and stress-related TFs (*FOS*, *JUN*, *JUNB*, *FOSB*, *CEBPD*, *BHLHE40*, *NFκB1*, *NFκB2* and *RELB*), decreased on D4 and D9 but increased on D11 and D16 (Figure 2F). A positive correlation between the hypoxia score and inflammatory response score was observed (Figure 2G), suggesting that hypoxia may be a key factor in regulating and activating inflammatory responses during mountaineering. Notably, the expression levels of genes related to the hypoxia pathway and *HIF1A* were significantly increased on D11 and D16 (Figure 2D and 2H). Hypoxia induces the upregulation of *HIF1A* expression, leading to the activation of HIF signaling and increased transcription of *NF-κB,* which serves as a downstream target of HIF[24], consistent with previous studies showing that *HIF1A* is essential for regulating the inflammatory response in myeloid cells[25].

### Dysregulation of pDCs in mountain climbers

The proportion of pDCs in mountain climbers exhibited a striking reduction on D16 (Figure 2C). pDCs can produce type I interferons (IFNs), secrete cytokines and chemokines, and capture and present antigens, thereby playing an important role in immunity to infections[26]. We investigated the changes in the molecular features of pDCs during high-altitude mountaineering, and DEGs and pathway enrichment analysis revealed that the downregulated DEGs were enriched mainly in the following pathways: ‘viral process’, ‘antigen processing and presentation’, ‘response to unfolded protein’, and ‘oxidative phosphorylation’ (Figure S6A). The downregulated DEGs during mountaineering mainly included the IFN-ɑ regulatory genes *IRF4* and *IRF7*, antigen presentation-related human leukocyte antigen (HLA) class II genes (*CD74*, *HLA-DRB5*, *HLA-DRB1*, *HLA-DRA*, *HLA-DQB1*, *HLA-DRP1* and *HLA-DPA1*), and the chemokine receptors *CXCR3* and *CXCR4* (Figure S6B). *CXCR3* and *CXCR4* play essential roles in the migration of pDCs from peripheral blood to infection sites[26]. In addition, we found that multiple DEGs related to protein secretion, such as S*EC61B*, *LMAN2*, *SSR3*, and *PDIA6*, were markedly downregulated (Figure S6B). Therefore, the observed reduction in the proportion of pDCs accompanied by the decrease in the expression level of key genes suggested that pDCs may experience functional impairment during high-altitude mountaineering.

Next, we explored the changes in the regulon activity of TFs in pDCs using SCENIC. The regulon activities of multiple TFs, such as inflammation- and stress-related TFs (*JUND*, *FOS* and *KLF6*), the IFN-related TF *IRF7* and the unfolded protein response (UPR)-related TFs *XBP1* and *ATF4*, began to decrease on D9 (Figure S6C). *IRF7* is a major transcriptional regulator of the production of type I IFNs in pDCs. *XBP1* is essential for pDC development and survival, and a lack of *XBP1* markedly reduces the size of the pDC compartment[27]. It was speculated that hypoxia may be a negative regulator of pDCs, which is consistent with previous studies showing that the proportion of pDCs significantly decreased under hypoxic conditions[10, 28]. Marked downregulation of the hypoxia pathway was detected in pDCs during mountaineering (Figure 2D). In particular, compared with other monocytes and cDCs, pDCs showed lower expression of *HIF1A* on D11 and D16 (Figure 2H), and in vitro studies have shown the essential role of *HIF1A* in type I IFN production by DCs[29]. Taken together, our findings indicate that mountain climbers may experience pDC dysregulation.

### Dynamic characterization of T and NK cell subsets during high-altitude mountaineering

T cell- and NK-cell-mediated immunity is one of the core immune functions, and we explored their functional changes during mountaineering. All CD8^+^ T, CD4^+^ T and NK cells were subclustered and classified into 20 subpopulations based on the expression of classical marker genes, including nine subgroups of CD8^+^ T cells and nonclassical T cells (n=135,616): naive CD8^+^ T cells, CD8^+^ effector memory T cells (CD8^+^ Tems), CD8^+^ central memory T cells (CD8^+^ Tcms), cytotoxic CD8^+^ T cells, terminal effector-like CD8^+^ T cells (TE-like CD8^+^ T cells), γδT cells, MAIT cells, proliferating T cells and NK-like T cells (Figure 3A and 3B); six CD4^+^ T cell subgroups (n=79,847): naive CD4^+^ T cells, IFN^+^ CD4^+^ T cells, CD4^+^ central memory T cells (CD4^+^ Tcms), T helper 2 cells (Th2 cells), Th17-like T cells and CD4^+^ regulatory T cells (Tregs) (Figure S7A and S7B); and five NK cell subgroups (n=69,673): CD56^+^ NK cells, active CD16^+^ NK cells, CD16^+^ NK cells, ZEB2^+^ NK cells and type 1 innate lymphoid cells (ILC1s) (Figure S7C and S7D). The abundances of the T and NK cell subgroups exhibited significant alterations during different periods of mountaineering (Figure 3C and S8A-S8D). The proportion of Tregs, the major suppressors of immune responses, was significantly increased on D9 (Figure 3C), and Tregs play crucial roles in the functional homeostasis of the immune system[30]. The proportion of naive CD4^+^ T cells decreased on D11, and that of Th2 cells increased on D11 (Figure 3C and S8D). Pathway analysis revealed that the T cell activation pathway was markedly activated in naive CD4^+^ T and Th2 cells on D11 and D16 (Figure S7E). These results suggested that Th2 cells may be activated during extreme altitude mountaineering.

**Figure 3.**
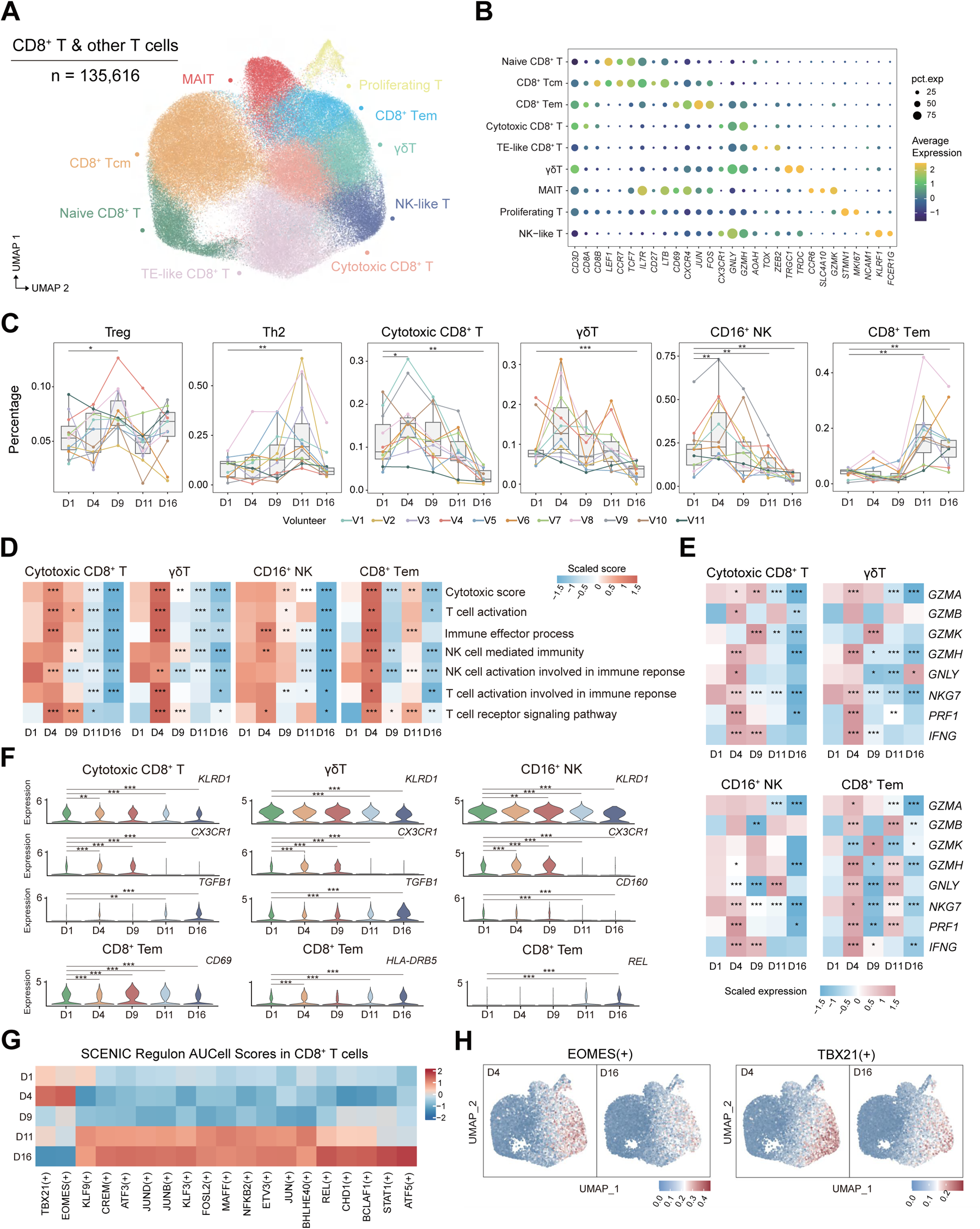
Dynamic changes in the functional status of T and NK cells in mountain climbers. (A) UMAP plot showing nine subsets of CD8^+^ T cells and other cells. (B) Dot plots depicting the percentages and average expression of the canonical genes in nine subsets of CD8^+^ T cells and other cells. (C) Boxplots showing the relative proportions of selected T and NK cell subsets during mountaineering (day 1, day 4, day 7, day 11 and day 16, n = 11); sample points from the same mountain climber are connected with the same color. The two-sided *p* values from the paired Wilcoxon rank-sum test are shown, \**p* < 0.05, \*\**p* < 0.01, \*\*\**p* < 0.001. (D) Heatmap showing the scaled scores for important immune-related pathways in cytotoxic CD8^+^ T, γδT, CD16^+^ NK and CD8^+^ Tem cells at different time points. The two-sided *p* values from the Wilcoxon rank-sum test are shown, \**p* < 0.05, \*\**p* < 0.01, \*\*\**p* < 0.001. (E) Heatmap showing the expression levels of cytotoxicity-related genes in cytotoxic CD8^+^ T, γδT, CD16^+^ NK and CD8^+^ Tem cells at different time points. The two-sided *p* values from the Wilcoxon rank-sum test are shown, \**p* < 0.05, \*\**p* < 0.01, \*\*\**p* < 0.001. (F) Violin plots showing dynamic changes in the expression of selected DEGs in cytotoxic CD8^+^ T, γδT, CD16^+^ NK and CD8^+^ Tem cells (|log_2_FC| ≥ 0.7, adjusted *p* < 0.05). \**p* < 0.05, \*\**p* < 0.01, \*\*\**p* < 0.001. (G) Heatmap showing the regulon AUCell scores for CD8^+^ T cells across the five time points, calculated using the SCENIC algorithm. (H) UMAP plots showing the regulon activities of *EOMES* and *TBX21* on D4 and D16 in CD8^+^ T cells, color-coded by AUCell score.

Notably, the proportions of cytotoxic CD8^+^ T, γδT and CD16^+^ NK cells exhibited similar variation trends: the proportion of cytotoxic CD8+ T cells increased on D4 but maintained a decreasing trend afterward and decreased to a minimum on D16 (Figure 3C). To investigate the dynamic changes in the functionality of these T and NK cells during different mountaineering stages, we identified DEGs (Table S5) and performed pathway analysis (Figure 3D). Overall, the pathway scores related to cytotoxicity, T cell activation, the immune effector process and the T cell receptor signaling pathway increased on D4 but significantly decreased on D11 and D16 (Figure 3D). Notably, the expression levels of cytotoxic genes (*GZMA*, *GZMK*, *GZMH* and *NKG7*) were also significantly reduced in cytotoxic CD8^+^ T, γδT and CD16^+^ NK cells on D11 and D16 (Figure 3E). Furthermore, the expression of *KLRD1,* which mediates inhibitory receptor signaling in T and NK cells, was significantly downregulated on D11 and D16 in cytotoxic CD8^+^ T, γδT and CD16^+^ NK cells (Figure 3F). Previous reports demonstrated that the deficiency of CD94 encoded by *KLRD1* resulted in susceptibility to lethal viral disease[31]. The expression of the chemokine receptor *CX3CR1*, which is expressed on cytotoxic T cells and NK cells and is thought to participate in T cell migration and immune memory formation[32], was significantly upregulated on D4 and D9 and downregulated on D11 and D16 in cytotoxic CD8^+^ T, γδT and CD16^+^ NK cells (Figure 3F). The expression of *CD160*, an activating receptor expressed by NK cells and a functional regulator of cytokine production by NK cells[33, 34], was significantly downregulated in CD16^+^ NK cells on D11 and D16 (Figure 3F). We also observed significant upregulation of the expression of the immunoregulatory gene *TGFB1* in cytotoxic CD8^+^ T cells and γδT cells on D11 and D16 (Figure 3F); this gene inhibits the differentiation and activation of T cells[35]. Overall, we observed enhanced phenotypes of cytotoxic CD8^+^ T, γδT and CD16^+^ NK cells on D4 during the period of acclimatization. However, on D11 and D16 during extreme altitude mountaineering, these cell populations exhibited altered phenotypes characterized by the downregulated expression of effector process-associated genes and the upregulated expression of the immunosuppressive molecule *TGFB1*.

Additionally, we identified activated CD8^+^ Tems that exhibited increased expression levels of activation-associated transcripts such as *CD69*, *CXCR4* and *JUN* (Figure 3B). Notably, the proportion of CD8^+^ Tems was significantly increased on D11 and D16 (Figure 3C). An elevated proportion of CD8^+^ CD69^+^ T cells has been previously reported following high-altitude trekking and is considered a beneficial response to immune perturbation[17]. To further investigate the transcriptional changes in CD8^+^ Tems during mountaineering, DEG and pathway analyses were performed. The pathway scores associated with T cell activation and cytotoxicity were increased on D4 but were significantly decreased on D11 in CD8^+^ Tems (Figure 3D). The expression of *CD69*, a classical marker of T cell activation, was significantly increased on D9 and D11 (Figure 3F). Along with the potential activation of CD8+ Tems, the expression of the cytotoxic serine protease *GZMB*, the antigen presentation-related HLA II class molecule *HLA-DRB5* and the *NF-κB* subunit *REL* was significantly upregulated on D11 and D16 in CD8^+^ Tems (Figure 3E and 3F).

Next, we analyzed the changes in TF regulon activity in CD8^+^ T and NK cells at different stages using SCENIC. The regulon activities of a set of TFs associated with inflammation and stress, including *JUN*, *REL*, *ATF3*, *ATF5* and *BHLHE40*, were obviously decreased on D4 and D9 but enhanced on D11 and D16 (Figure 3G). Notably, the regulon activities of *TBX21* and *EOMES* increased on D4 and then obviously decreased, especially on D16 (Figure 3G and 3H). *TBX21* and *EOMES* are highly expressed in CD8^+^ T and NK cells[36] and regulate the expression of downstream genes, such as *GZMA*, *CCL5*, *NKG7*, *CD160* and *SPON2* (Figure S7F). The regulon activities of *TBX21* and *EOMES* may be suppressed during extreme altitude mountaineering. In general, these results suggested that mountain climbers may experience impaired functionality in cytotoxic CD8^+^ T, γδT and CD16^+^ NK cells on D11 and D16.

### Enhanced unfolded protein response in plasma B cells during high-altitude mountaineering

B cells play an important role in humoral immunity. We subclustered 54,546 B cells and annotated them into eight subgroups based on the expression of classical marker genes, including six B-cell subsets (naive B cells, IFN^+^ B cells, activated B cells, memory B cells, FCRL3^+^ B cells and SOX5^+^ B cells) and two plasma cell subsets (plasma B cells and proliferating plasma B cells) (Figure 4A and 4B). Then, we analyzed the compositional changes in different stages and observed a significant reduction in the proportion of naive B cells on D4, accompanied by a significant increase in activated B cells. The plasma B cells and other B cell subgroups did not exhibit statistically significant changes (Figure 4C, S9A and S9B). To reveal the functional changes in B cells in mountain climbers, we performed DEG analysis (Table S6). The expression of HLA class II genes (*HLA-DRB5* and *HLA-DQA2*) in activated B cells significantly increased during mountaineering, indicating an augmentation of B cell-mediated antigen presentation capabilities (Figure 4D). In addition to the marked increase in the expression level of *CD69*, we also discovered a significant upregulation of *CD83* on D11 and D16 (Figure 4D), which is important for B cell activation[37].

**Figure 4.**
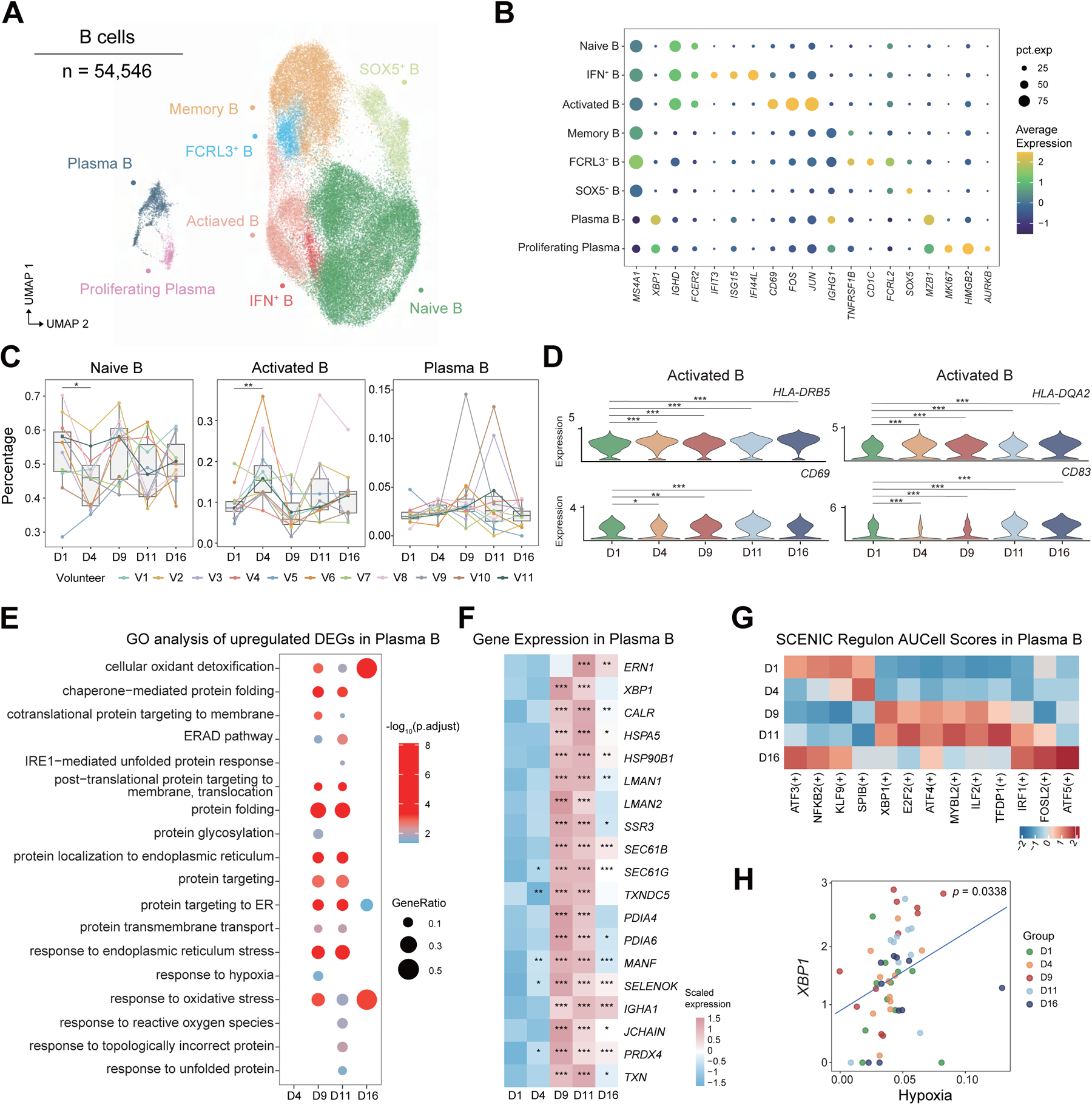
Activation of the unfolded protein response in plasma B cells during mountaineering. (A) UMAP plot showing eight B cell subsets. (B) Dot plots depicting the percentages and average expression levels of the canonical genes in eight subsets of B cells. (C) Boxplots showing the relative proportions of selected B cell subsets during mountaineering (day 1, day, day 7, day 11 and day 16, n = 11); sample points from the same mountain climber are connected with the same color. The two-sided *p* values from the paired Wilcoxon rank-sum test are shown, \**p* < 0.05, \*\**p* < 0.01, \*\*\**p* < 0.001. (D) Violin plots showing dynamic changes in the expression of selected DEGs in activated B cells(|log_2_FC|≥0.7, adjusted *p* < 0.05). \**p* < 0.05, \*\**p* < 0.01, \*\*\**p* < 0.001. (E) The fifteen enriched biological processes according to Gene Ontology (GO) analysis of the upregulated DEGs in plasma B cells. Dot color indicates the statistical significance of the enrichment, and dot size represents the gene ratio annotated to each term. (F) Heatmap illustrating scaled expression values of selected DEGs across time points in plasma B cells (|log_2_FC| ≥ 0.7, adjusted *p* < 0.05). \**p* < 0.05, \*\**p* < 0.01, \*\*\**p* < 0.001. (G) Heatmap showing the regulon AUCell scores from plasma B cells across different time points, calculated using the SCENIC algorithm. (H) Correlation analysis between hypoxia scores and *XBP1* expression levels in plasma B cells for each climber all five time points (linear mixed model).

To further investigate the functional changes in plasma B cells during mountaineering, DEG analysis was performed (Table S6). Pathway enrichment analysis revealed that the upregulated DEGs were mainly enriched in pathways related to protein folding and the UPR (Figure 4E). The UPR pathway is an essential component of plasma B cell differentiation, and plasma B cells employ a specialized UPR dependent on the IRE1-XBP1 axis for protein synthesis[38]. The expression of *ERN1* (also known as IRE1), a major sensor of the UPR, was significantly upregulated on D11 and D16 (Figure 4F). The expression of *XBP1* significantly increased on D9 and D11 (Figure 4F). Additionally, we observed prominently upregulated expression of protein secretion-related genes (*LMAN1*, *HSPA5*, *SSR3* and *PDIA4*) and the immunoglobulin genes *IGHA1* and *JCHAIN* (Figure 4F). The activation of the UPR in plasma B cells potentially contributes to the enhancement of the antibody secretion response in mountain climbers, which is consistent with the reported increase in immunoglobulin levels at high altitudes[39].

The development of plasma B cells requires the involvement of specific TFs. We also identified *XBP1* in plasma B cells using SCENIC, and the regulon activity of *XBP1* was upregulated on D9 and D11 (Figure 4G). As a critical member of the UPR, *XBP1* regulates the terminal differentiation of B cells into plasma B cells[40] and controls downstream genes involved in protein synthesis (Figure S9C). Hypoxia can induce *XBP1* mRNA expression and activate its mRNA splicing[41], and a positive correlation between the hypoxia score and the expression of *XBP1* was observed in plasma B cells (Figure 4H). Taken together, these results demonstrated that high-altitude mountaineering induced the activation of the UPR in plasma B cells, which may contribute to the enhancement of humoral immunity in mountain climbers.

### Metabolic reprogramming of circulating immune cells in mountain climbers

Pathway enrichment analysis revealed a reduction in the levels of oxidative phosphorylation in CCL3^+^ CD14^+^ Monos and pDCs during mountaineering (Figure S5C and S6A). Considering the profound impact of metabolic programs on immune cell function[18], an analysis of metabolic features centered on oxidative phosphorylation and the glycolysis pathway across different cell types was performed. The expression level of oxidative phosphorylation increased on D4 but decreased significantly on D9, D11 and D16, especially in monocytes (Figure 5A and 5B). In contrast, the expression level of glycolysis decreased on D4 and D9 but significantly increased on D11 and D16 (Figure 5A). This change was consistent with the observed variation in the hypoxia pathway, which was initially suppressed and subsequently activated (Figure 2D and S10A). A positive correlation between the hypoxia score and glycolysis score was observed in different immune cell subgroups (Figure S11A and S11B), suggesting that hypoxia may be a key factor in regulating the metabolic program of immune cells during mountaineering. The expression of mitochondrial electron transport chain (ETC) subunits was downregulated. For instance, the expression of *ATP5F1E* (ATP synthase complex) was significantly downregulated during mountaineering in CCL3^+^ CD14^+^ Monos, FCN1^+^ CD14^+^ Monos, C1^+^ CD16^+^ Monos and cDC2s (Figure 5C). Decreased expression levels of *COX4I1*, *COX7A2*, *COX5B* and *COX7C* (complex IV) were observed in IFN^+^ CD16^+^ Monos (Figure 5D). It was evident that the activity of the mitochondrial ETC decreased under high-altitude hypoxic conditions. HIF is widely recognized as the primary regulator of metabolic adaptation to hypoxia and can drive a switch from oxidative to glycolytic metabolism during hypoxia[42]. Interestingly, significantly upregulated *HIF1A* expression was observed on D11 and D16 in most subgroups, except for pDCs (Figure 2H and S10A). Moreover, the expression of *LDHA* (lactate dehydrogenase A) and *GAPDH* (glyceraldehyde 3-phosphate dehydrogenase) was significantly elevated during mountaineering in CD8^+^ Tems and plasma B cells, respectively (Figure 5E). *LDHA* encodes a key enzyme in glycolysis that facilitates the conversion of pyruvate to lactate. *GAPDH* is an essential component of glycolysis that catalyzes the conversion of glyceraldehyde-3-phosphate to 1,3-bisphosphoglycerate[43]. We also observed upregulated expression of the glucose transporter *SLC2A3* on D11 and D16(Figure 5E); this gene facilitates cellular glucose uptake to support the glycolytic pathway[44]. Taken together, these observations suggested that the circulating immune cells of mountain climbers may undergo metabolic reprogramming.

**Figure 5.**
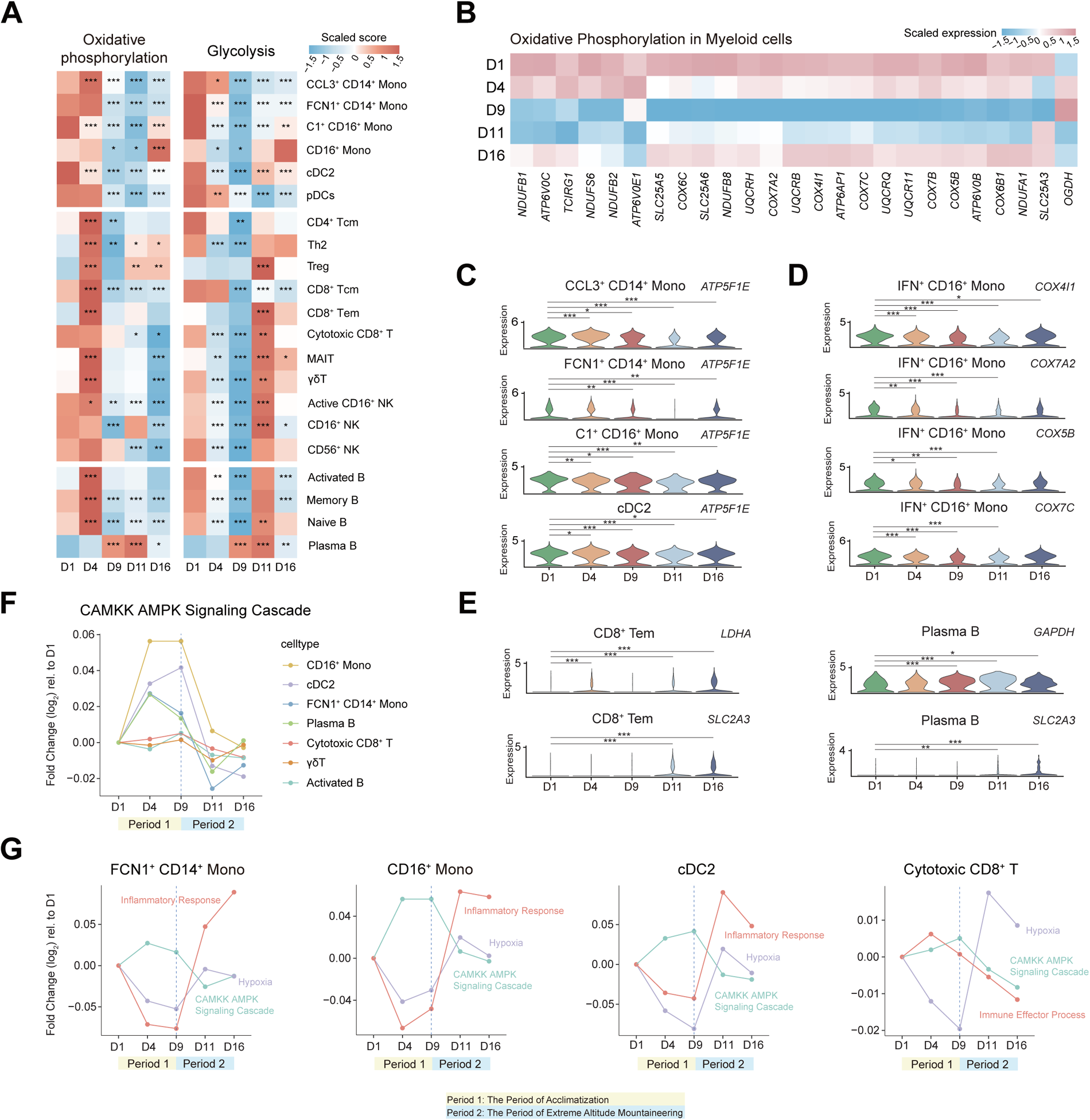
Metabolic profile of circulating immune cells during mountaineering. (A) Heatmap showing the scaled scores for oxidative phosphorylation and glycolysis pathways across different time points in selected cell subsets. The two-sided *p* values from the Wilcoxon rank-sum test are shown, \**p* < 0.05, \*\**p* < 0.01, \*\*\**p* < 0.001. (B) Heatmap showing the average expression of selected genes from oxidative phosphorylation pathways in myeloid cells across different time points. (C) Violin plots showing dynamic changes in the expression of *ATP5F1E* in CCL3^+^ CD14^+^ Monos, FCN1^+^ CD14^+^ Monos, C1^+^ CD16^+^ Monos and cDC2s (|log_2_FC| ≥ 0.7, adjusted *p* < 0.05). \**p* < 0.05, \*\**p* < 0.01, \*\*\**p* < 0.001. (D) Violin plots showing dynamic changes in the expression of *COX4I1*, *COX7A2*, *COX5B* and *COX7C* in IFN^+^ CD16^+^ Monos (|log_2_FC| ≥ 0.7, adjusted *p* < 0.05). \**p* < 0.05, \*\**p* < 0.01, \*\*\**p* < 0.001. (E) Violin plots showing dynamic changes in the expression of glycolysis-related genes in plasma B cells and CD8^+^ Tems (|log_2_FC| ≥ 0.7, adjusted *p* < 0.05). \**p* < 0.05, \*\**p* < 0.01, \*\*\**p* < 0.001. (F) Dynamic changes in AMPK-related pathways (‘CAMKK AMPK Signaling Cascade’) during high-altitude mountaineering in different immune cell subsets. Stage 1 is the acclimatization period, and stage 2 is the extreme altitude mountaineering period. Dots represent the mean log_2_-fold change relative to baseline (D1). (G) The biological processes that significantly changed during high-altitude mountaineering in FCN1^+^ CD14^+^ Monos, CD16^+^ Monos, cDC2s and cytotoxic CD8^+^ T cells included the AMPK signaling, inflammatory response, hypoxia and immune effector process pathways. Dots represent the mean log_2_-fold change relative to baseline (D1).

Cell energy sensors also play important roles in the metabolic regulation of immune cells[15]. AMP-activated protein kinase (AMPK) serves as the main regulator of intracellular energy homeostasis and can be activated by various stimuli, including cellular stress and physical activity[45]. The activity of the AMPK-related pathway ‘CAMKK AMPK Signaling Cascade’ was significantly increased on D4 and D9, particularly in monocytes; however, it decreased on D11 and D16 (Figure 5F and S10B). The expression of the AMPK subunits *PRKAB1* and *PRKAB2*, as well as the upstream kinase *CAMMK2*, was upregulated on D4 and D9 in CCL3^+^ CD14^+^ Monos, FCN1^+^ CD14^+^ Monos, cytotoxic CD8^+^ T cells and γδT cells (Figure S10C). In addition, the expression level of *PRKAA1* was also significantly upregulated on D4 and D9 in cytotoxic CD8^+^ T cells (Figure S10C). *PRKAA1* is crucial for the proliferation and differentiation of cytotoxic T lymphocytes[46]. Activation of the AMPK pathway can not only promote the metabolic adaptation of immune cells but also exert a wide variety of immunoregulatory effects[47]. For example, the AMPK pathway exerts inhibitory effects on the inflammatory response of macrophages and DCs while promoting T cell effector function[48, 49]. In our study, we observed that during the period of acclimation, when the AMPK-related pathway was activated, the inflammatory response in monocytes and cDC2s was suppressed, and the immune effector process of cytotoxic CD8^+^ T cells was upregulated (Figure 5G). In contrast, the activity of the AMPK-related pathway decreased during extreme altitude mountaineering, whereas the activity of the hypoxia pathway increased, accompanied by the activation of inflammatory responses in monocytes and cDC2s and a decrease in the immune effector process in cytotoxic CD8^+^ T cells (Figure 5G). Overall, our findings suggested that the dynamic activation of the AMPK and hypoxia pathways induced by high-altitude mountaineering may play crucial roles in regulating the metabolism and function of immune cells.

### The defense of circulating immune cells against oxidative stress in mountain climbers

Previous findings have indicated that exercise at high altitudes increases the generation of ROS[17]. When ROS production exceeds the load capacity of the antioxidant system, it can lead to oxidative stress. The pathway scores for ‘Cellular Response to Reactive Oxygen Species’ and ‘Cellular Response to Oxidative Stress’ were calculated for each immune subgroup, and we found that the scores decreased on D4 and D9 and then increased on D11 and D16 in most subgroups, in addition to pDCs and several B cell subgroups (Figure 6A). A positive correlation between the hypoxia score and oxidative stress score was observed for circulating immune cells (Figure 6B). These findings indicated that circulating immune cells manifested distinct reactions to oxidative stress during high-altitude mountaineering.

**Figure 6.**
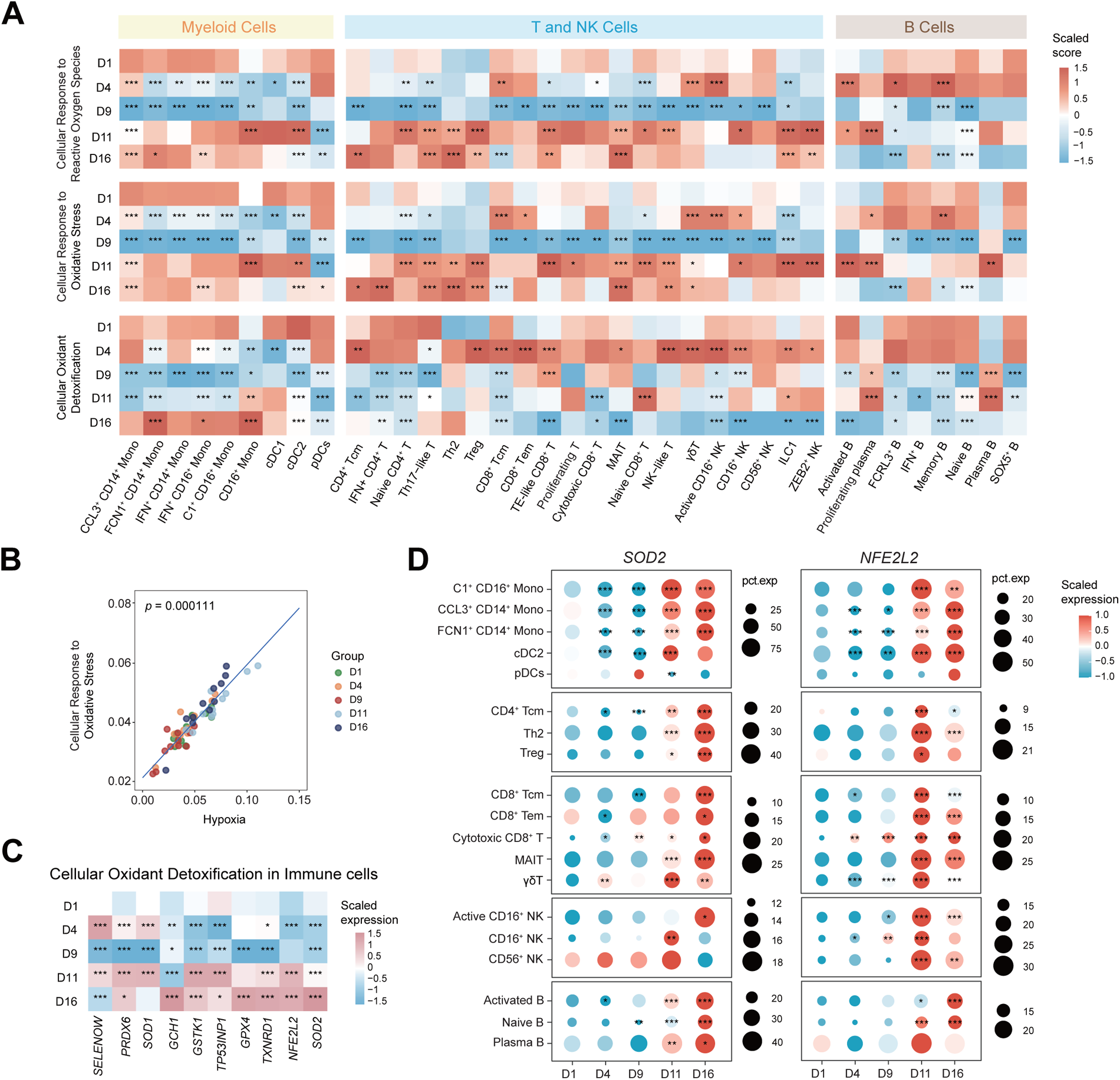
Oxidative stress response of circulating immune cells during mountaineering. (A) Heatmap showing the scaled scores for reactive oxygen species, oxidative stress and oxidant detoxification-related pathways across different time points. The two-sided *p* values from the Wilcoxon rank-sum test are shown, \**p* < 0.05, \*\**p* < 0.01, \*\*\**p* < 0.001. (B) Correlation analysis between hypoxia scores and oxidative stress scores for each climber at all five time points (linear mixed model). (C) Heatmap showing the scaled expression of selected genes from cellular oxidant detoxification pathways in circulating immune cells across different time points. The two-sided *p* values from the Wilcoxon rank-sum test are shown, \**p* < 0.05, \*\**p* < 0.01, \*\*\**p* < 0.001. (D) Dot plots showing the scaled average expression levels and percentages of *SOD2* and *NFE2L2* in selected immune cell subsets at different time points. The two-sided *p* values from the Wilcoxon rank-sum test are shown, \**p* < 0.05, \*\**p* < 0.01, \*\*\**p* < 0.001.

Moreover, cells are equipped with an antioxidant system that scavenges excess ROS to reduce oxidative damage[50]. We next calculated the pathway scores for ‘Cellular Oxidant Detoxification’ in different immune subsets and observed increased scores in T and NK cells on D4, whereas the scores were generally lower in most subgroups on D9, D11 and D16 (Figure 6A). Notably, the oxidative detoxification scores were greater on D11 and D16 in monocytes and plasma B cells (Figure 6A). The expression of multiple antioxidant genes in different mountaineering stages, including *SOD2*, *NFE2L2*, *SOD1*, *GPX4* and *TXNRD1*, was significantly upregulated on D11 and D16 (Figure 6C). Among them, *SOD2* and *NFE2L2* levels were significantly increased in monocytes and DCs but not in pDCs (Figure 6D). The SOD2 enzyme is specifically localized to the mitochondrial matrix at the main site of superoxide production and catalyzes the reaction of superoxide to less reactive hydrogen peroxide[51]. *NFE2L2* (also known as *NRF2*) serves as a central regulator that responds to oxidative stress by controlling a battery of defensive genes encoding antioxidant and detoxification enzymes[52]. In addition, we observed that the expression of *PRDX4* and *TXN* was significantly elevated in plasma B cells during mountaineering (Figure 4F). In contrast, markedly decreased expression of *SOD1*, *TXN*, *PRDX5* and *PRDX6* was observed in pDCs (Figure S6B). *TXN* and *PRDXs* are major members of the thioredoxin system, which is one of the central antioxidant systems and plays a vital role in maintaining the intracellular ROS balance[53]. In particular, *PRDX4*, a typical endoplasmic reticulum (ER)-resident antioxidant that scavenges excess H_2_O_2_, is essential for the correct folding of proteins[54]. Generally, when the level of oxidative stress increased on D11 and D16, monocytes and plasma B cells exhibited enhanced oxidative detoxification ability, while pDCs and most T and NK cells exhibited decreased oxidative detoxification ability. These results highlighted the importance of the enhanced antioxidant capacity of circulating immune cells to defend against elevated oxidative stress and further maintain their immune functions.

### Plasma metabolic changes in mountain climbers

Metabolic reprogramming in immune cells is observed during mountaineering, and metabolic changes in immune cells may be localized, but if they occur across multiple cells or tissues, such changes can result in systemic metabolic alterations[18]. To explore the plasma metabolic alterations in mountain climbers during mountaineering, we conducted VIP and c-means clustering analysis. According to the VIP scores, the levels of some metabolites, including 3-methyloxindole, N-acetyl-L-aspartic acid, asparagine, glycine and palmitoylcarnitine, increased during mountaineering, while the levels of other metabolites, such as glucose and piperine, decreased (Figure 7A). Four change patterns were identified in different periods of mountaineering: cluster 1 contained analytes whose levels were sharply decreased on D4 and D9, followed by analytes whose levels were increased on D11 and decreased on D16; cluster 4 exhibited the opposite trend (Figure 7B); the analytes in cluster 2 exhibited a sharp decrease on D4 and subsequently remained at a low level; and the analytes in cluster 3 showed a pattern of a gradually decreasing trend (Figure 7B). The differentially abundant metabolites were mainly distributed in cluster 1 and cluster 4, whereas differential lipids exhibited a predominant distribution in cluster 2 and cluster 3 (Figure 7B).

**Figure 7.**
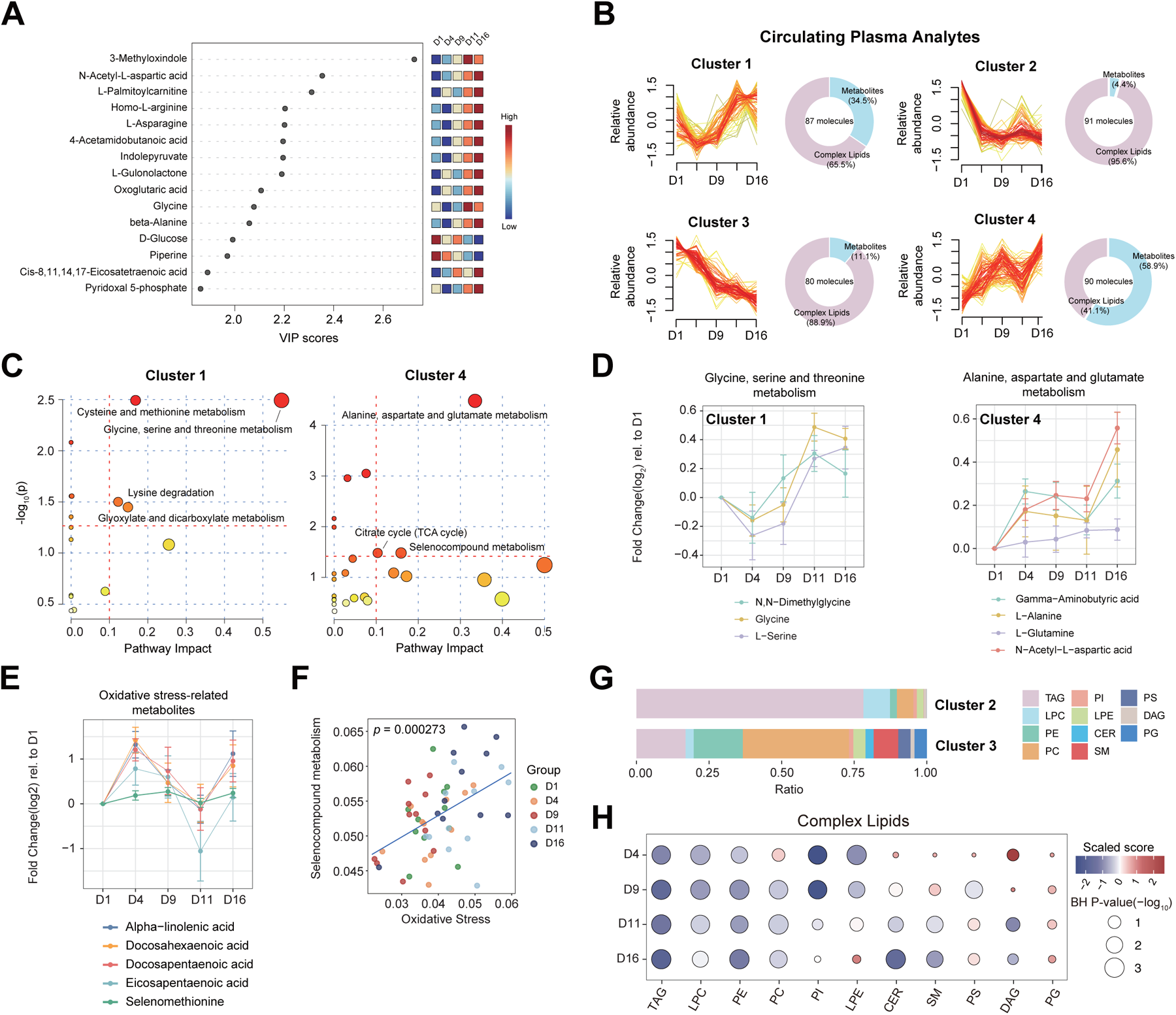
Time series analysis of plasma metabolomics and lipidomic data during high-altitude mountaineering. (A) Variable importance in projection (VIP) scores showing the top 15 differentially abundant metabolites for each sampling point. The heatmap on the right displays the scaled value of the metabolite at each time point. (B) Clustering of differentially abundant metabolites and lipids. The circular plots on the right show the percentage of differentially abundant metabolites and lipids for each cluster. (C) Pathway enrichment analysis of metabolites from cluster 1 and cluster 4. The red horizontal dashed line indicates the threshold of statistical significance for the y-axis calculated by the hypergeometric test (*p* < 0.05). The red vertical dashed lines indicate pathway impact values ≥ 0.1. (D) The dynamic changes in key metabolites in the enrichment pathways of cluster 1 and cluster 4. Dots represent the mean log_2_-fold change relative to D1, and horizontal bars represent the standard error of the mean (SEM). (E) Dynamic changes in oxidative stress-related metabolites during mountaineering. The dots represent the mean log_2_-fold change relative to day 1, and the bars represent the standard error of the mean (SEM). (F) Correlation analysis between oxidative stress scores and selenocompound metabolism scores for each climber at all five time points (linear mixed model). (G) Stacked column charts showing the percentage of each complex lipid in cluster 2 and cluster 3. (H) Chemical classification enrichment analysis of complex lipids. The pathway direction is the change in significant molecules in each classification relative to D1 (blue, downregulated; red, upregulated). The size of the dot represents the significance of the pathway.

The analytes in cluster 1 were mainly enriched in serine and glycine metabolism, representative metabolites that included glycine, serine and methionine (Figure 7C and S12A). The levels of serine and glycine decreased on D4 and D9, followed by an increase on D11 and D16 (Figure 7D). The concentration of plasma glycine is regulated by glucagon, which stimulates the degradation of glycine, and glucagon deficiency elevates the level of glycine[55]. Furthermore, intense physical activity may result in increased muscle protein breakdown and the subsequent release of amino acids into the bloodstream[56], potentially resulting in increased glycine levels. The increased availability of glycine can be utilized for the synthesis of serine and glutathione. The role of glutathione as a vital antioxidant lies in its ability to reduce cellular damage induced by oxidative stress[57]. The analytes in cluster 2 were not enriched in metabolic pathways, and those in cluster 3 were mainly involved in tryptophan metabolism (Figure S12B).

The analytes in cluster 4 were enriched in glutamine metabolism, and representative metabolites included glutamine, N-acetyl-L-aspartic acid, alanine and gamma-aminobutyric acid (GABA) (Figure 7C, 7D and S12A). Glutamine is the most abundant circulating amino acid in plasma, as it is a carbon source second only to glucose, which can generate 2-oxoglutarate through glutaminolysis[58]. Consistently, the levels of citrate cycle (TCA cycle)-related substances, including 2-oxoglutarate and isocitric acid, increased during mountaineering (Figure S12C). It has been suggested that mountain climbers may rely on glutamine as an important anaplerotic substrate to complete the TCA cycle during mountaineering, consistent with previous studies showing that high altitude-exposed individuals presented increased plasma glutamine[59]. Importantly, glutamine can actively improve immune function, and glutamine supplementation may reduce the incidence of infections reported by marathon runners[60]. In addition, GABA, a neurotransmitter, has a variety of biological functions, including antifatigue and immune regulation activities[61].

Analytes in cluster 4 were also involved the metabolism of selenocompounds, from which selenomethionine (Se-Met) (Figure 7C and S12C), a natural organic selenium, is a key metabolite. As the main source of selenium in humans, Se-Met has been shown to have important antioxidant properties involving the formation of antioxidant enzymes[62]. Se-Met increased on D4 and D9 but significantly decreased on D11 (Figure 7E). The decreased level of Se-Met may be related to the increased oxidative stress response observed in immune cells, as a significant positive correlation between selenocompound metabolism and oxidative stress in circulating immune cells was observed (Figure 7F). Intriguingly, we observed a similar variation trend in omega-3 polyunsaturated fatty acids (n-3 PUFAs), alpha-linolenic acid, docosahexaenoic acid, docosapentaenoic acid and eicosapentaenoic acid (Figure 7E). PUFAs are vulnerable to ROS attack due to their structural instability, resulting in a marked increase in the end-product malondialdehyde (MDA), which is a biomarker of oxidative stress[63].

In addition, fatty acid oxidation may be activated on D11 and D16, as reflected by the accumulation of acylcarnitines and medium- to long-chain fatty acids (Figure S12D). Acylcarnitine is an intermediate product of fatty acid oxidation, and the heart is considered the main contributor to the increased abundance of long-chain acylcarnitines in plasma[64], such as palmitoylcarnitine and stearoylcarnitine (Figure S12D). These changes were concomitant with the accumulation of 3-hydroxybutyric acid, the main ketone body in the blood (Figure S12E). Ketone bodies are predominantly produced from fatty acid β-oxidation-derived acetyl-CoA in the liver and serve as key contributors to energy metabolism in the context of energy-depleted states[65]. These observations suggested that the utilization of fatty acids might be enhanced during extreme altitude mountaineering.

Clusters 2 and 3 mainly contained complex lipids, such as triacylglycerol (TAG), lysophosphatidylcholine (LPC), phosphatidylethanolamine (PE), phosphatidylcholine (PC), phosphatidylinositol (PI), lysophosphatidylethanolamine (LPE), ceramide (CER) and sphingomyelin (SM) (Figure 7B and 7G). The role of lipids is critical for cell structure, intracellular and extracellular signaling and energy storage. The levels of TAG, LPC, PE and PC decreased significantly during mountaineering (Figure 7H). TAG molecules represent the major form of fatty acid storage within cells, and in the plasma, prolonged exercise has been reported to accelerate plasma TAG clearance[66]. LPC can activate a variety of signaling pathways in monocytes/macrophages through G protein-coupled receptors and Toll-like receptors, further increasing the production of proinflammatory cytokines, whose downregulated expression is considered associated with worse health status[67]. PE and PC are the most abundant phospholipids in mammalian cell membranes and play crucial roles in regulating lipid metabolism and maintaining membrane protein stability[68]. The progressive decrease in plasma PE levels during mountaineering may be regarded as a potentially positive change[69]. The levels of PI and LPE were significantly decreased on D4 and D9 and then gradually recovered, while the levels of CER and SM were significantly decreased on D4 and D9 (Figure 7H). CER is the precursor of most SM. The plasma concentration of SM is altered in various metabolic disorders, serving as a valuable indicator for prognosis and diagnosis[70]. Overall, the observed decrease in lipid levels may reflect lipid metabolism disorders during mountaineering.

## DISCUSSION

Extreme adventure sports, such as high-altitude mountaineering, have become increasingly popular. High-altitude mountaineering involves exposure to a hypobaric hypoxic environment, leading to a series of physiological, metabolic and immunological changes. Circulating immune cells, including monocytes, DCs, T cells, NK cells and B cells, function as “sentinel tissue” to monitor the body’s response[71]. Previous studies have proposed peripheral blood lymphocytes as biosensors for whole-body responses to high altitudes[16]. However, few studies have investigated the response profiles of circulating immune cells during mountaineering. We identified and characterized the immune response and metabolic changes during high-altitude mountaineering. Importantly, the mountaineering plan consisted of two stages (the first being the period of acclimatization, and the second being the period of extreme altitude mountaineering). Sampling at multiple time points enabled a more comprehensive understanding of the immune response alterations and adaptation characteristics of mountain climbers in different periods.

Research in the field of exercise immunology has demonstrated that physical activity can induce significant alterations in the immune response, contingent upon exercise intensity, duration, and type[72]. Moderate-intensity exercise is acknowledged to confer a multitude of immunological benefits, including augmented immunosurveillance, diminished systemic inflammation levels, and heightened lymphocyte concentrations, whereas prolonged or high-intensity exercise is associated with immune dysfunction, elevated inflammation levels and heightened susceptibility to infectious disease[7, 8]. Our analysis revealed that high-altitude mountaineering induced intricate and dynamic immune alterations in mountain climbers. Climbers exhibited distinct immune alterations in the two stages of mountaineering. During the acclimation period, for myeloid cells, we observed the significant suppression of the inflammatory response in monocytes and cDCs. The immunosuppressive phenotype of CCL3^+^ CD14^+^ Monos was subsequently characterized by notable the upregulation of *S100A8* and *S100A9* expression and marked downregulation of the expression of multiple inflammatory genes on D4 and D9. The upregulation of *S100A8* expression has been documented in individuals acutely exposed to high altitudes[73]. For lymphocytes, the proportions of cytotoxic CD8^+^ T, γδT and CD16^+^ NK cells were significantly increased on D4, and the expression of effector process-related genes was significantly upregulated on D4. Notably, AMPK-associated pathways were also activated on D4 and D9 in monocytes, cDC2s, cytotoxic CD8^+^ T cells and γδT cells.

As one of the central regulators of cellular metabolism, AMPK plays a critical role in orchestrating cellular metabolic reprogramming. Under conditions of low energy availability, AMPK phosphorylates downstream targets to enhance adenosine triphosphate (ATP) production while reducing ATP consumption[45]. Previous studies have demonstrated that exercise-induced activation of AMPK can confer a wide range of health benefits, making it an attractive target for the treatment of immune-related diseases[47, 74]. In myeloid cells, AMPK exerts inhibitory effects on inflammatory responses[48], highlighting the intricate interplay between inflammation and AMPK signaling. In T cells, AMPK plays a pivotal role in regulating T cell-mediated adaptive immunity[49]. This finding aligned with our observation that during the acclimation period, when the AMPK-related pathway was activated, there was a concomitant suppression of the inflammatory response in monocytes and cDCs and an increase in effector process scores for cytotoxic CD8^+^ T cells.

During extreme high-altitude mountaineering, monocyte- and cDC-mediated inflammatory responses were significantly activated on D11 and D16, with increased expression of *IL1B*, *TNF* and *NFκB1*, in line with the observed increase in CRP levels during trekking in the Himalayas[5]. Proinflammatory *IL1B* expression has been reported to be correlated with worsening symptoms of acute mountain sickness (AMS)[75], and the exacerbation of the inflammatory response in high-altitude hypoxic environments may contribute to the pathogenesis of diseases, such as chronic obstructive pulmonary disease or chronic obstructive sleep apnea[76]. Compared to those on D1, the proportions of cytotoxic CD8^+^ T, γδT and CD16^+^ NK cells on D16 were significantly lower, accompanied by the downregulated expression of genes associated with immune effects, including *GZMA*, *GZMK*, *GZMH*, *NKG7*, *KLRD1*, *CD160* and *CX3CR1*. These findings are consistent with the observed decrease in the percentage of cytotoxic T cells following a marathon[20]. The impairment of T cell-mediated function at extreme altitudes might disrupt immune homeostasis and increase the risk of infectious disease. These observations may be attributed to the increased intensity of physical activity during extreme altitude mountaineering (D11 and D16), which could further exacerbate the degree of hypoxia. Importantly, the activity of the hypoxia pathway was significantly activated on D11 and D16 in most immune cells.

The effect of hypoxia on the inflammatory response has been widely reported in studies of high-altitude environments[24]. On the one hand, hypoxia-induced *HIF1A* can upregulate the expression of *NF-κB*; on the other hand, hypoxia promotes the release of *NF-κB* and its translocation to the nucleus by activating the IKKβ pathway, further upregulating the transcription of inflammatory genes[77]. Among T and NK cells, the upregulation of the immunoregulatory gene *TGFB1* was observed in cytotoxic CD8^+^ T and γδT cells on D11 and D16. Hypoxia-induced *TGFB1* may exert inhibitory effects on the expression of *TBX21* and *EOMES* in T cells[35, 78]. Similarly, *TGFB1* also downregulated the expression of *TBX21* in NK cells[79]. Within plasma B cells, hypoxia may promote the activity of *XBP1*[41], which controls the downstream genes involved in protein synthesis. Notably, the activity of the hypoxia pathway and the expression of *HIF1*A were significantly increased in most cell types during extreme altitude mountaineering, while decreased hypoxia scores and low *HIF1A* expression were observed in functionally impaired cells, such as pDCs, confirming the key role of hypoxia in regulating the immune response of mountain climbers.

Furthermore, another intriguing finding in our study is the adaptation characteristics of circulating immune cells during extreme altitude mountaineering. Considering the exposure to high-altitude hypoxic conditions coupled with intense physical activity, the energy and metabolic homeostasis of mountain climbers may be challenged substantially. We observed a decrease in oxidative phosphorylation activity and an increase in glycolytic activity on D11 and D16 in most immune cells. Previous studies have reported that climbers experience attenuated mitochondrial function after exposure to high altitudes[16, 80]; these findings imply that the functionality of mitochondria might be constrained under high-altitude conditions. Notably, the expression of the glycolysis-related genes *LDHA*, *GAPDH* and *SLC2A3* was significantly enhanced on D11 and D16 in CD8^+^ Tems and plasma B cells. HIF is known to play an important role in hypoxia-induced reprogramming, and increased *HIF1A* expression was observed on D11 and D16 in most immune cells. Moreover, we observed increased pathway activity in most subgroups in response to ROS and oxidative stress on D11 and D16, which was consistent with previous studies showing that exercise at high altitudes increased the levels of oxidative stress[13]. In the context of increased oxidative stress on D11 and D16, monocytes and plasma B cells showed increased expression of antioxidant-related genes (*NFE2L2*, *SOD2*, *PRDX4* and *TXN*) on D11 and D16. *NFE2L2* is the major transcription regulator in the cellular response to oxidative stress[52]. An increase in SOD activity is considered a crucial adaptive mechanism for coping with hypoxic conditions at high altitudes to reduce oxidative damage[81].

Finally, the high-resolution analysis of plasma analytes revealed numerous molecules affected by high-altitude mountaineering, and these metabolic processes were involved in energy metabolism (amino acid and fatty acid metabolism) and oxidative stress (selenocompound metabolism), suggesting the crucial involvement of metabolites in metabolic adaptation in climbers. In high-altitude hypoxic environments, mountain climbers are in a low-sugar state, which may improve their energy supply by promoting glutamine metabolism and fatty acid metabolism. Interplay among metabolism, oxidative stress and immunity was observed, reflected in selenocompound metabolism, and a positive correlation between selenocompound metabolism and oxidative stress was also observed in circulating immune cells.

In conclusion, the results of our study provide a multi-omics profile of the response to high-altitude mountaineering. We used a single-cell transcriptome to analyze the circulating immune profile of climbers, revealed distinct immune responses during two stages of mountaineering, and explored the adaptive characteristics of circulating immune cells during extreme altitude mountaineering. The enhanced expression of glycolysis and antioxidant genes was shown to play a crucial role in supporting the functionality of immune cells. Conversely, decreased expression of those genes was observed in pDCs with impaired immune function. Furthermore, the energy status of mountain climbers can potentially be promoted through the enhancement of glutamine metabolism and fatty acid metabolism.

## Data availability

The scRNA-seq data generated by this study were deposited at the China National GeneBank (CNGB) Sequence Archive (https://db.cngb.org/cnsa/) under accession nos. Other relevant data are available from the corresponding author upon reasonable request.

## Code availability

R codes and other custom scripts are available upon reasonable request.

## Acknowledgements

We sincerely thank the eleven mountain climbers who enthusiastically participated in this mountaineering expedition. This work was supported by the Guangzhou Basic and Applied Basic Research Program (202201010189) and the National Key R&D Program of China (2022YFC3400400). We would also like to acknowledge the support from the Chinese National Gene Bank.

## Author contributions

Experimental strategy and design: J.W., X.J., C.Y.L., J.C., L.Q.L., J.H.Y.

Laboratory experiments: Y.W., X.M.L., Y.T.H., Y.Y., Y.L.H., C.L., G.X.H., H.R.T., H.H.L., Y.Z., G.D.Z.

Statistical analyses: S.C.Y., J.Z.L., Z.L.H., X.W., Y.L., W.K.W., Y.H.Z., C.W.

Manuscript writing: J.H.Y., J.Z.L., S.C.Y.

Manuscript editing: J.H.Y., J.Z.L., W.W.Z., R.Q.L.,

All authors critically revised the manuscript for important intellectual content and gave final approval for the version to be published. All authors agree to be accountable for all aspects of the work in ensuring that questions related to the accuracy or integrity of any part of the work are appropriately investigated and resolved.

## METHODS

### Sample collection and ethics statement

This study was performed during an expedition to summit the Muztagh Ata peak (7,546 m), and eleven healthy volunteers residing at low altitudes (< 300 m) were recruited to participate in the expedition. None of the individuals had known chronic diseases, such as cardiovascular or respiratory system disorders. During mountaineering, we monitored the dynamic changes in 10 physiological parameters (e.g., routine blood tests, SpO_2_, HR, LLS and body composition) of climbers using professional equipment. To investigate alterations in metabolism and the immune response during high-altitude mountaineering, 55 blood samples were collected from climbers at five distinct time points. The research protocol received approval from BGI’s institutional review board of bioethics and biosafety, and written informed consent was obtained from each individual participating in the mountaineering expedition.

### Protocol for the mountaineering expedition

For the mountaineering itinerary, participants reached 1,289 m on the first day (D1), and the mountaineering expedition was divided into two phases: an initial acclimatization period (D2-D8) followed by an extreme altitude mountaineering period (D9-D15). The participants embarked on a trek to reach 3,200 m and subsequently arrived at the base camp (4,400 m) on D3. From there, acclimatization to high altitudes was carried out at the base camp. During this period, the participants ascended to Camp 1 (5,500 m) on D5, followed by a two-day rest at base camp before proceeding to Camp 2 (6,300 m) on D9. Finally, the participants undertook a four-day consecutive ascent from base camp to the summit (7,546 m). Before reaching the summit, five out of the eleven individuals utilized supplemental oxygen for a duration of time on D15. On D16, the participants returned 1,289 m. During the expedition, 55 blood samples at five different time points (D1, D4, D9, D11 and D16) were collected. Fasting blood samples for each participant were collected at 9:00 a.m.; the fifth sampling was performed before dinner due to itinerary constraints. No medications (including acetazolamide) were taken by the participants during the expedition. According to the LLS, none of the eleven participants suffered from severe acute mountain sickness. For additional details, see Supplementary Table 1.

### Plasma and peripheral blood mononuclear cells (PBMCs) separation

Plasma and PBMC isolation were conducted with the following protocols. The whole blood of the mountain climbers was collected in 5mL EDTA anticoagulant tubes. Briefly, 5 mL of whole blood was centrifuged at 940×g for 10 min, after which the upper plasma layer was transferred to a 15-mL centrifuge tube, collected after centrifugation at 2,600×g for 10 min, aliquoted, and stored at - 80 °C. The lower immune cells were diluted with 1x phosphate-buffered saline (PBS) supplemented with 2% bovine serum albumin (BSA) and transferred to SepMate^TM^–50 tubes. After centrifugation at 1200×g for 10 min, the PBMC layer was poured into a new centrifuge tube. The cells were washed twice with 3 mL of 1× PBS containing 2% fetal bovine serum (FBS) and centrifuged at 300×g for 8 min. The supernatant was discarded, and the PBMC pellet was resuspended in FBS supplemented with 10% dimethylsulfoxide (DMSO). Afterwards, the frozen tubes were transported on dry ice to be stored in liquid nitrogen at the end of the expedition.

### Preparation of single-cell suspensions

To thaw the frozen PBMCs, the cells were placed in a 37 °C water bath for 2 min, after which the cells were gently poured into a 15-mL centrifuge tube. The thawed cells were resuspended in 10 mL of preheated 1× PBS supplemented with 10% FBS at 37 °C and mixed gently. After centrifugation at 500×g for 5 min at room temperature, the supernatant was removed, and the cell pellet was resuspended in 2 mL of PBS supplemented with 0.04% BSA. The cells were strained through a 40-μm cell strainer, and the filtered cells were collected and then resuspended at a concentration of 800-1,500 cells/μL. The cell suspensions were maintained on ice for sequencing.

### Sequencing library construction

The sequencing libraries were constructed according to the instructions of the DNA Nanoball (DNB) elab C4 scRNA Preparation Kit (MGI). The prepared single-cell suspension, the barcode mRNA capture beads, and the droplet generation oil were loaded into the corresponding reservoirs on a microfluidic chip for droplet generation. The cells were lysed, and the released mRNA was captured and barcoded in the droplets. Following reverse transcription, complementary DNA (cDNA) was generated and amplified. The amplified cDNA was quantified using a BioAnalyzer High Sensitivity Chip (Agilent) and subsequently fragmented to 400-600 bp with NEBNext dsDNA Fragmentase. The cDNA and oligo library construction steps were performed as follows. The libraries were made of DNBs and sequenced on an ultrahigh-throughput DNBSEQ-T1 sequencer at the CNGB. cDNA libraries were generated using the following read lengths: 41-bp read 1 (containing a 20-bp cell barcode and a 10-bp unique molecular identifier (UMI)), 100-bp read 2, and a 10-bp sample index. The following oligo libraries were used: 26-bp read 1, 42-bp read 2, and 10-bp sample index.

### Single-cell RNA-seq data processing

Sequencing data were processed using the Open Source Pipeline (https://github.com/MGI-tech-bioinformatics/DNBelab_C_Series_HT_scRNA-analysis-software). First, bead barcodes and UMI sequences were extracted using the parsing function in PISA (https://github.com/MGI-tech-bioinformatics/DNBelab_C_Series_HT_scRNA-analysis-software/software/PISA). Reads with incorrect barcodes were excluded based on the use of a barcode whitelist. The scRNA-seq data were then aligned to the GRCh38 (hg38) reference genome using STAR[82]. Finally, gene expression levels in cells were assessed using PISA, and gene-cell matrices were created for each library.

Then, we used the R package Seurat (version 4.4.0) to apply preliminary counts to downstream data analysis, including quality control, normalization, feature selection, dimensionality reduction, unsupervised clustering, and visualization[83]. First, cells with more than 10% mitochondrial genes, more than 1% erythrocyte genes, and less than 200 or more than 6,000 genes were excluded. Then, for each library, DoubletFinder (version 2.0.3) was used to remove doublets[84]. The top 5% of cells most similar to the “pseudo-doublet” were excluded. Finally, genes expressed in fewer than three cells, as well as mitochondrial genes, noncoding RNA genes, and ribosomal genes were filtered. In addition, a portion of the cells were clustered together by red blood cells contamination with highly expressed red blood cell genes, and we removed the red blood cell clusters to eliminate the effect on clustering. Subsequently, the “NormalizeData” function was used to convert the data into logarithmic space, and the “FindVariableGene” function was used to find the top 2,000 variable genes in the dataset. Then, “ScaleData” was used to remove unwanted variations due to differences in sequencing depth. The first 20 principal components (PCs) were computed using RunPCA. Finally, Harmony (version 1.1.0) was used to correct for batch effects of different library.

### Unsupervised clustering and subclustering

The Louvain algorithm was used as a modular optimization technique to set the granularity of the clusters at a resolution of 0.75. The nonlinear dimensionality reduction technique “UMAP” was used to visualize the clusters in two-dimensional space. Cellular identity was determined by identifying differentially expressed genes (DEGs) for each cluster using the “FindALLMarkers” function with default parameters and comparing those genes to known cell type-specific genes. The clusters were annotated based on typical marker expression, and 15 cell subpopulations were identified. After scaling, myeloid, CD4^+^ T, CD8^+^ T, NK, and B cell clusters were extracted as separate Seurat objects; the principal components were identified, each subset was clustered at a unique resolution using the Louvain algorithm, and the clusters were annotated based on dimensionality and gene expression associated with the phenotypes of the typical subsets described above.

### DEGs and GO enrichment analysis

To search for DEGs in each cell subgroup during mountaineering, we used the “FindMarkers” function (Seurat). For each cell subgroup, we computed the fold change in mean gene expression levels between D1 and subsequent time points, and the p value was calculated utilizing the Wilcoxon rank sum test. To annotate the functions of these DEGs, we performed GO pathway enrichment analysis by using the clusterProfiler package (version 4.10.0) in R (OrgDb = org.Hs.eg.db, pvalueCutoff = 0.05, qvalueCutoff = 0.05, pAdjustMethod = “BH”) to enrich the DEGs in GO terms[85]. For DEG details, see Supplementary Tables 4, 5 and 6.

### Module scores for feature gene set expression

To assess the dynamic changes in biological processes mediated by immune cell subsets, we employed the “AddModuleScore” (Seurat) function to assess the scores for the gene sets. Briefly, the score for each cell was defined as the average expression of the featured gene set minus the average expression of the corresponding control feature gene set. The referenced gene sets are listed in Supplementary Table 7 and were collected from the Molecular Signatures Database (MSigDB)[86]. The Wilcoxon rank-sum test was used to evaluate the statistical significance of differences between D1 and subsequent time points.

### Gene regulatory network analysis based on SCENIC

To predict the potential transcriptional regulatory networks in different immune cell subgroups, we used pySCENIC (version 0.12.1). Initially, we employed GRNBoost to infer all potential targets of transcription factors. Subsequently, the output coexpression module was utilized in conjunction with the cisTarget database to identify putative regulators. The regulon areas under the curve per cell (AUCell) module of pySCENIC was used to analyze regulator activity, and active regulators were identified using the default thresholds[23]. The differentially expressed regulators were identified using the R package limma (version 3.50.0) with the following parameters: adjusted *p* ≤ 0.05 and |logFoldChange| ≥ 0.4. For regulator details, see Supplementary Table 8.

### Target metabolite quantification by HM Meta Assays

The HM Meta Assay (BGI, Shenzhen, China) was utilized for quantifying metabolites. This assay covers a wide range of chemical classes, including bile acids, amino acids, fatty acids, carboxylic acids, hydroxyl acids, phenolic acids and indoles, by measuring 700 compounds. Calibration curves were generated using serial dilutions of metabolite standards. In 96-well plates, 20-μL aliquots of either standards or plasma samples were mixed with 120 μL of internal standard solution (methanol-diluted metabolite internal standards). The plates were then stored at -20 °C for 20 min, followed by centrifugation at 4,000 × g and 4 °C for 30 min. The supernatants (30 μL) were transferred to new plates for derivatization with freshly prepared reagents (10 μL), which included a mixture of 200 mM 3-NPH in 75% aqueous methanol and 96 mM EDC-6% pyridine in methanol. Derivatization took place at 30 °C for one hour. Prior to LC‒MS analysis, the samples were diluted with ice-cold 50% methanol and then transferred to new plates containing 135 μL of the supernatant.

A Waters ACQUITY UPLC instrument coupled to a SCIEX QTRAP 6500 PLUS mass spectrometer equipped with an ESI source controlled by Analyst software was used for UPLC-MRM analysis. Chromatographic separation was performed on a Waters ACQUITY BEH C18 column (1.7 μm particle size; dimensions: 100 mm × 2.1 mm). Both positive and negative ion modes were employed during instrument operation. Ultimately, HMQUANT software (BGI, Shenzhen, China) was used for the absolute quantification of metabolites.

### Target lipid quantification by HM lipid assays

Lipid measurement was conducted using the HM Lipid Assay (BGI, Shenzhen, China). This assay covers a wide range of lipid subclasses and quantifies 1600 lipid molecules, including 18 lipid classes: SM, CER, cholesterol ester (CE), monoacylglycerol (MAG), diacylglycerol (DAG), TAG, lysophosphatidic acid (LPA), phosphatidic acid (PA), LPC, PC, LPE, PE, lysophosphatidylinositol (LPI), PI, lysophosphatidylglycerol (LPG), phosphatidylglycerol (PG), lysophosphatidylserine (LPS) and phosphatidylserine (PS). Calibration curves were generated using serial dilutions of lipid standards. In 96-well plates, aliquots of either standards or plasma samples were mixed with internal standard solution (precooled isopropanol with SPLASH internal standards, avanti) (20 μL each). The plates were then stored at -20 °C for 20 min before being centrifuged at 4,000 × g and 4 °C for 30 min. Supernatants containing lipids were transferred to new plates for LC-MS analysis in negative mode, while a portion of the supernatants was diluted into methanol for LC-MS analysis in positive mode.

UPLC-MS/MS analysis was performed on a Waters ACQUITY UPLC coupled to a SCIEX QTRAP5500 PLUS mass spectrometer equipped with an ESI source controlled by Analyst software. Chromatographic separation was performed on a Waters ACQUITY CSH C18 column (1.7 μm, 100 mm × 2.1 mm). SM, CE, Cer, TAG, DAG, and MAG were quantified in positive mode, whereas the phospholipids and lysophospholipids were quantified in negative mode utilizing the scheduled MRM settings. HMQUANT software (BGI, Shenzhen, China) incorporating isotope correction via the LICAR package was used for absolute lipid quantification.

To ensure the accuracy of the analytical results, we discarded lipids detected in less than 2/3 of the samples, and missing values were estimated using the lower limit from the corresponding lipid[87].

### Screening for differential plasma metabolites and lipids

To identify differential plasma metabolites and lipids, we first calculated VIP scores using PLS-DA in the R package MetaboAnalystR (version 4.0)[88]. We analyzed metabolites with VIP scores ≥ 1 as differentially abundant metabolites and lipids for downstream data analysis. For details on the metabolites and lipids, see Supplementary Tables 2 and 3.

### Fuzzy c-means clustering

Fuzzy c-means clustering was performed using the R package Mfuzz (version 2.62.0)[89]. We selected the screened differentially abundant metabolites and lipids for clustering.

### Plasma metabolite pathway analysis

We performed pathway enrichment analysis of the metabolites obtained from fuzzy c-means clustering by group separately using the pathway analysis module in the MetaboAnalystR package. For bubble plots, we selected metabolic pathways with p values ≤ 0.05 and impacts ≥ 0.1[90].

### Linear mixed model

Since the assumption of independence cannot be met for repeated measurements, zone group data, and spatially correlated data, the above data are often analyzed using linear mixed-effects models (e.g., correlation analysis)[91]. The following model was implemented:

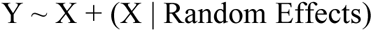

Mixed linear models include both fixed and random effects, where fixed effects are the variables under study and random effects are the influences that we wish to control for, which are referred to as “Groups” in this paper. The mixed linear model was generated using the R package statistical function lme4 (version 1.1-35.1). We tested the difference between the fixed-effects model and the model with fixed effects removed through ANOVA to determine whether the linear correlation between the fixed effects was significant.

### Quantification and statistical analysis

Statistical analysis was performed using R (version 4.3.1). The Wilcoxon rank-sum test was used in this study. Multiple comparison adjustments were performed using the Benjamini-Hochberg method. \**p* < 0.05, \*\**p* < 0.01, \*\*\**p* < 0.001.

## SUPPLEMENTAL INFORMATION

**Supplementary Figure 1.**
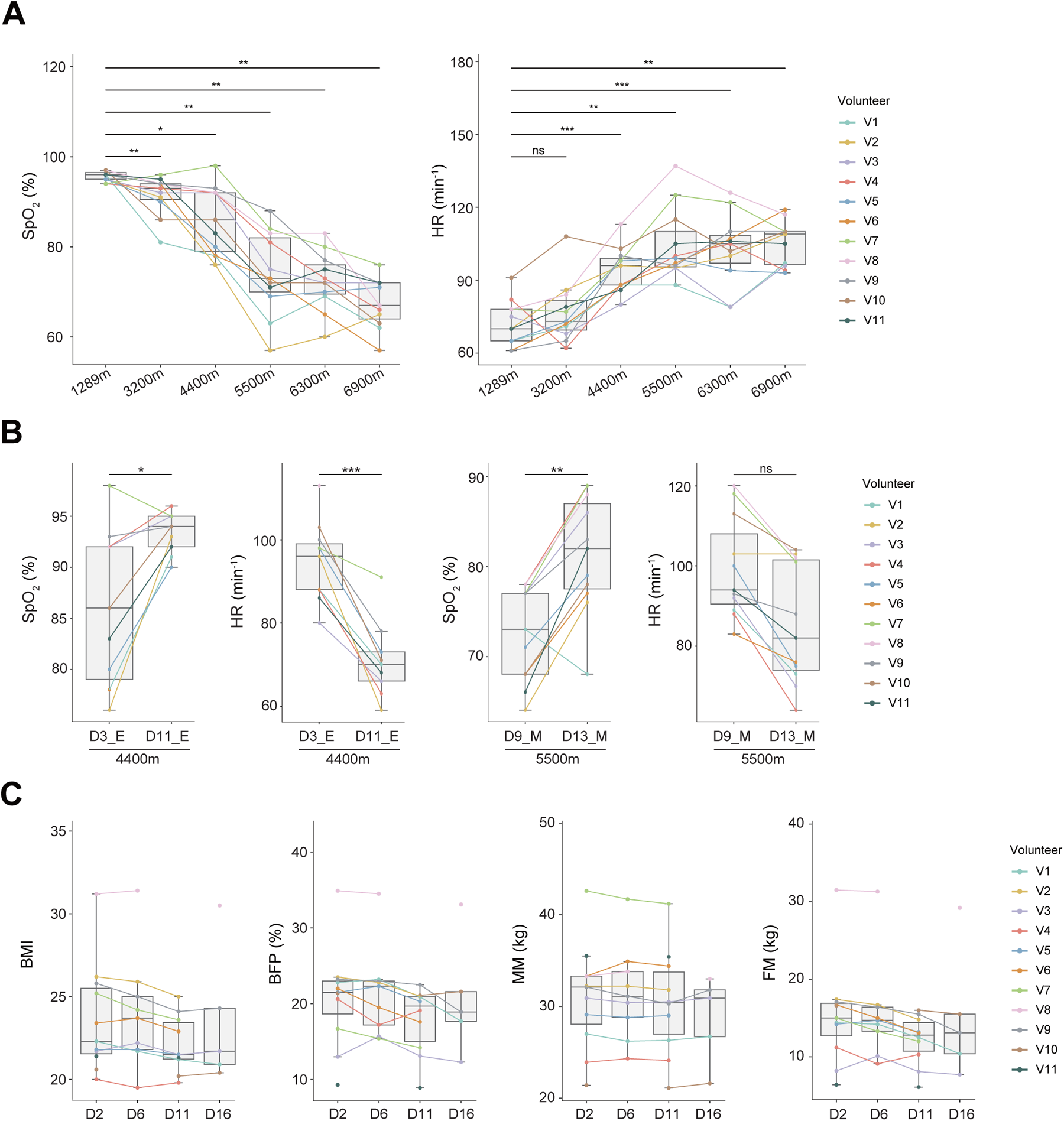
Effects of high altitude on physiological parameters during mountaineering. (A) Box plots showing the dynamic changes in SpO_2_ and heart ratHR with increasing altitude; sample points from the same mountain climber are connected with the same color (n = 11). The two-sided *p* values from the paired Wilcoxon rank-sum test are shown, \**p* < 0.05, \*\**p* < 0.01, \*\*\**p* < 0.001. (B) Box plots showing the changes in SpO_2_ and HR when re-exposed to the same altitude (4,400 m and 5,500 m); sample points from the same mountain climber are connected with the same color (n = 11). The two-sided *p* values from the paired Wilcoxon rank-sum test are shown, \**p* < 0.05, \*\**p* < 0.01, \*\*\**p* < 0.001. (C) Box plots showing the changes in body composition during mountaineering; sample points from the same mountain climber are connected with the same color. The two-sided *p* values from the paired Wilcoxon rank-sum test are shown, but no significant differences were detected. Body mass index (BMI), body fat percentage (BFP%), muscle mass (MM, kg) and fat mass (FM, kg) were measured.

**Supplementary Figure 2.**
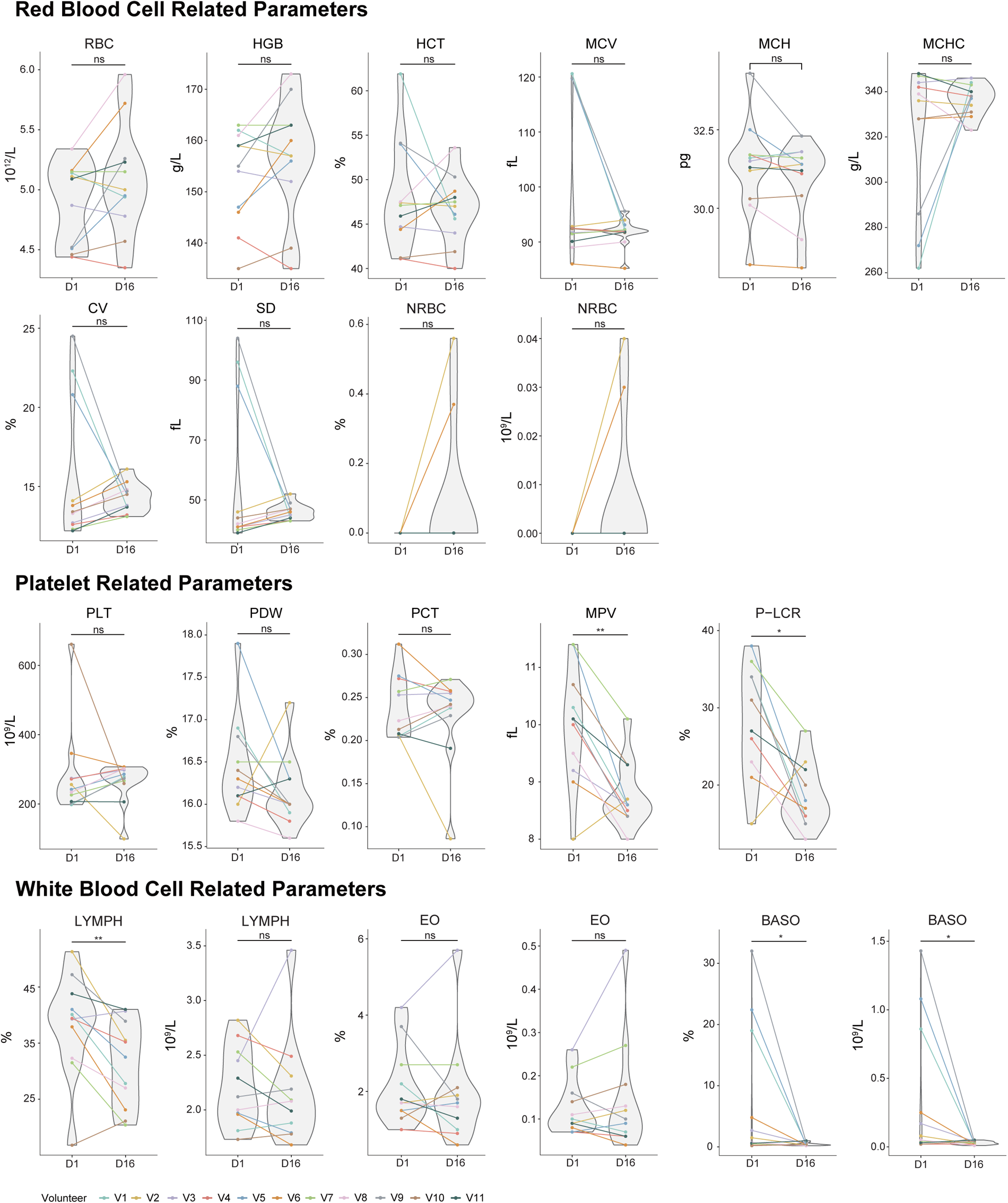
Effects of high altitude on routine blood parameters during mountaineering. Violin plots showing the changes in routine blood parameters before (D1) and after (D16) mountaineering, divided into three groups: red blood cell-related parameters, platelet-related parameters and white blood cell-related parameters. The complete names of these parameters are displayed in Table S1. The two-sided *p* values from the paired Wilcoxon rank-sum test are shown, \**p* < 0.05, \*\**p* < 0.01, \*\*\**p* < 0.001.

**Supplementary Figure 3.**
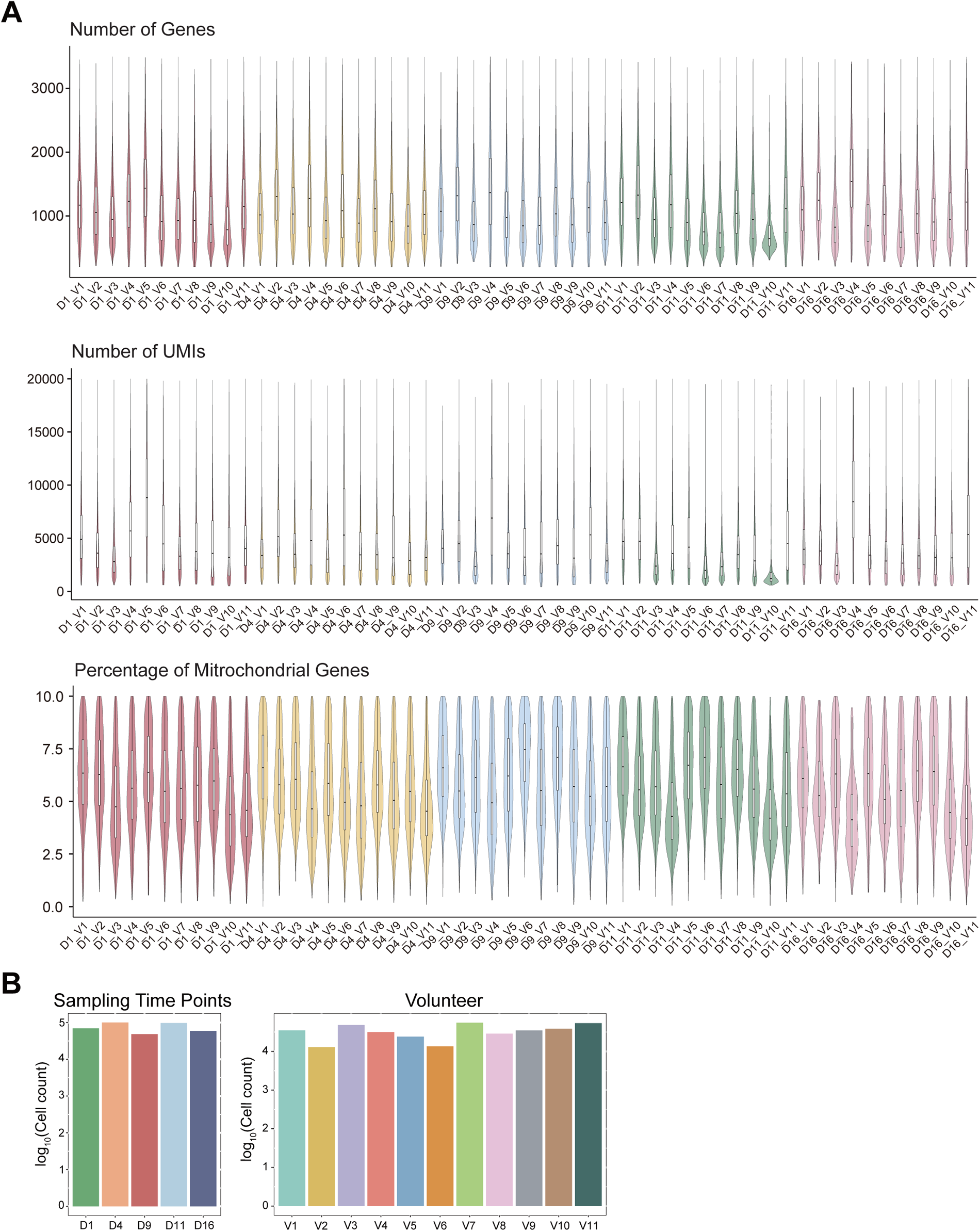
Quality control of the scRNA-seq data. (A) Violin plots showing the distributions of gene counts per cell, unique molecular identifier (UMI) per cell, and percentage of mitochondrial transcripts per cell across five time points from eleven mountain climbers. (B) Bar plots showing the number of cells at each time point and the number of cells from each volunteer. D1 (n = 69,847 cells), D4 (n = 100,271 cells), D9 (n = 48,686 cells), D11 (n = 97,447 cells), and D16 (n = 5,9471 cells).

**Supplementary Figure 4.**
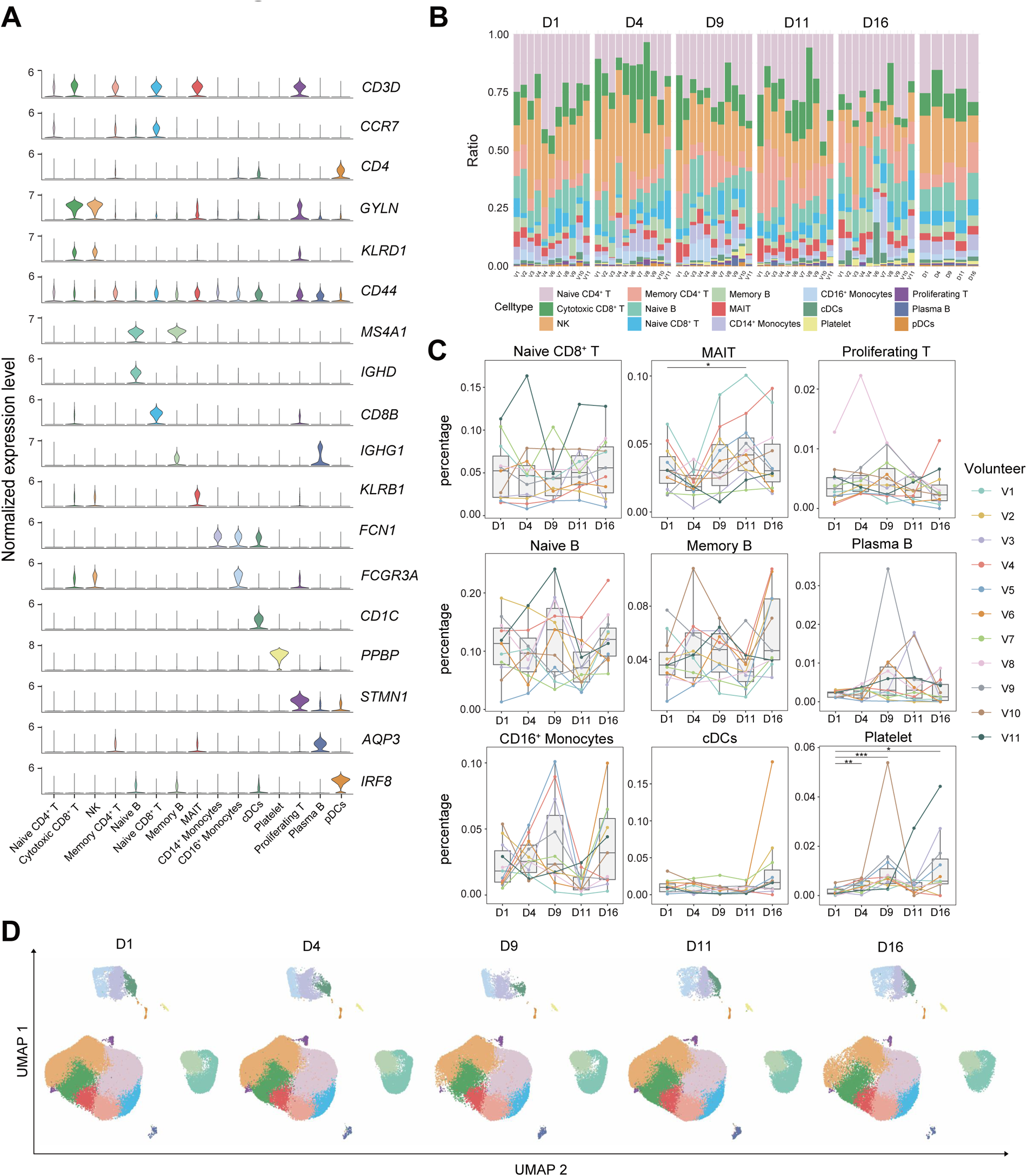
Single-cell analysis of circulating immune cells in mountain climbers. (A) Violin plots depicting the normalized expression levels of canonical markers associated with each main immune cell subgroup. (B) Stacked column charts showing the proportions of each cell type from each volunteer and different sampling time points. (C) Boxplots showing the relative proportions of immune cell subsets during mountaineering (day 1, day 4, day 7, day 11 and day 16, n = 11); sample points from the same mountain climber are connected with the same color. The two-sided *p* values from the paired Wilcoxon rank-sum test are shown, \**p* < 0.05, \*\**p* < 0.01, \*\*\**p* < 0.001. (D) UMAP plot of all immune cell subtypes identified from single-cell samples from climbers at different time points, colored according to their respective subgroup.

**Supplementary Figure 5.**
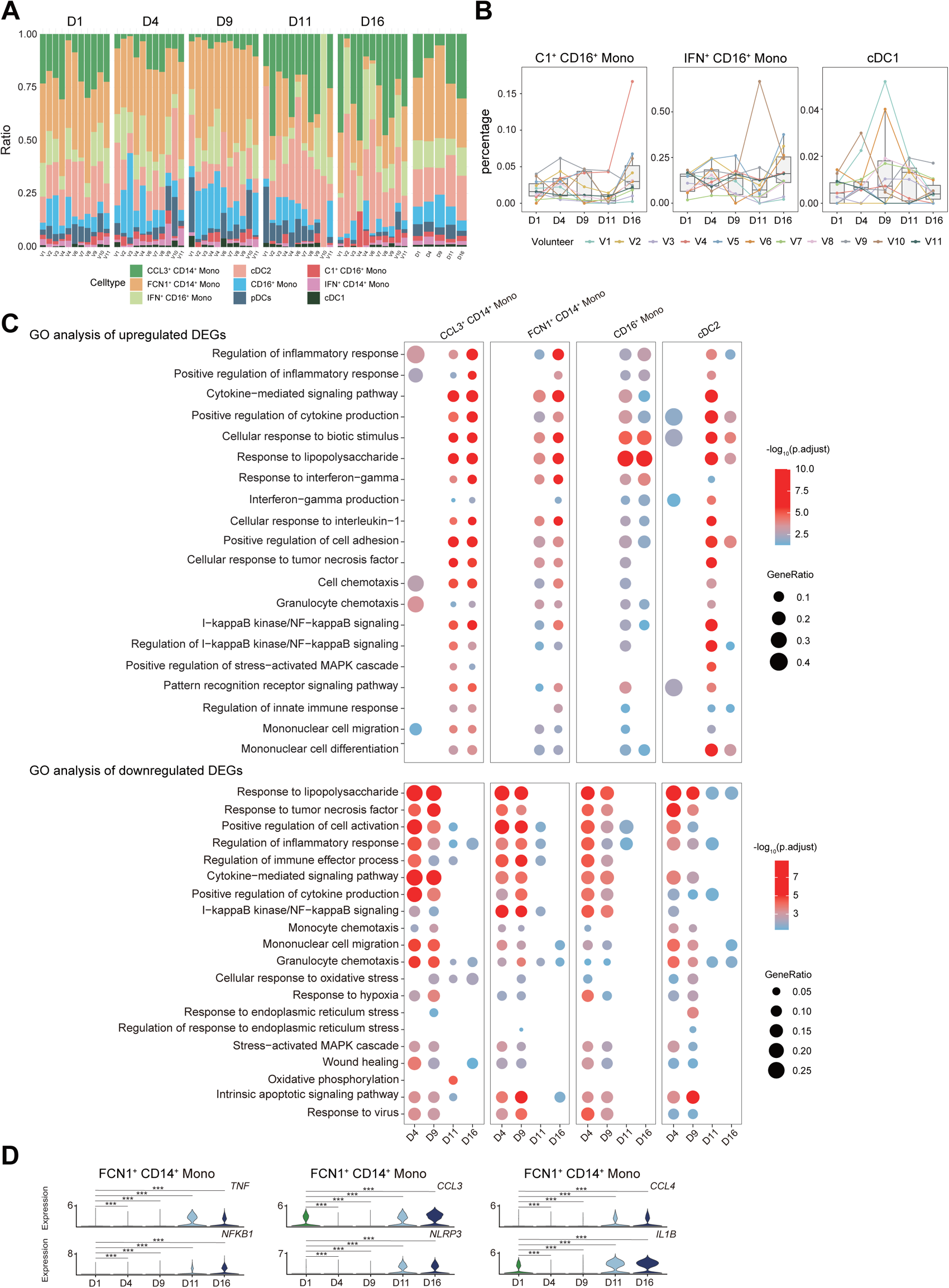
Transcriptional changes in monocyte subsets during mountaineering. (A) Stacked column charts showing the proportions of myeloid cell subsets from each volunteer and different sampling time points. (B) Boxplots showing the relative proportions of myeloid cell subsets during mountaineering (day 1, day 4, day 7, day 11 and day 16, n = 11); sample points from the same mountain climber are connected with the same color. The two-sided *p* values from the paired Wilcoxon rank-sum test are shown, \**p* < 0.05, \*\**p* < 0.01, \*\*\**p* < 0.001. (C) Dot plots showing the enriched biological processes by GO analysis of upregulated and downregulated DEGs on day 4, day 7, day 11 and day 16 compared to day 1. Dot color indicates the statistical significance of the enrichment, and dot size represents the gene ratio annotated to each term. (D) Violin plots showing dynamic changes in the expression of selected DEGs in FCN1^+^ CD14^+^ Monos (|log_2_FC| ≥ 0.7, adjusted *p* < 0.05). \**p* < 0.05, \*\**p* < 0.01, \*\*\**p* < 0.001.

**Supplementary Figure 6.**
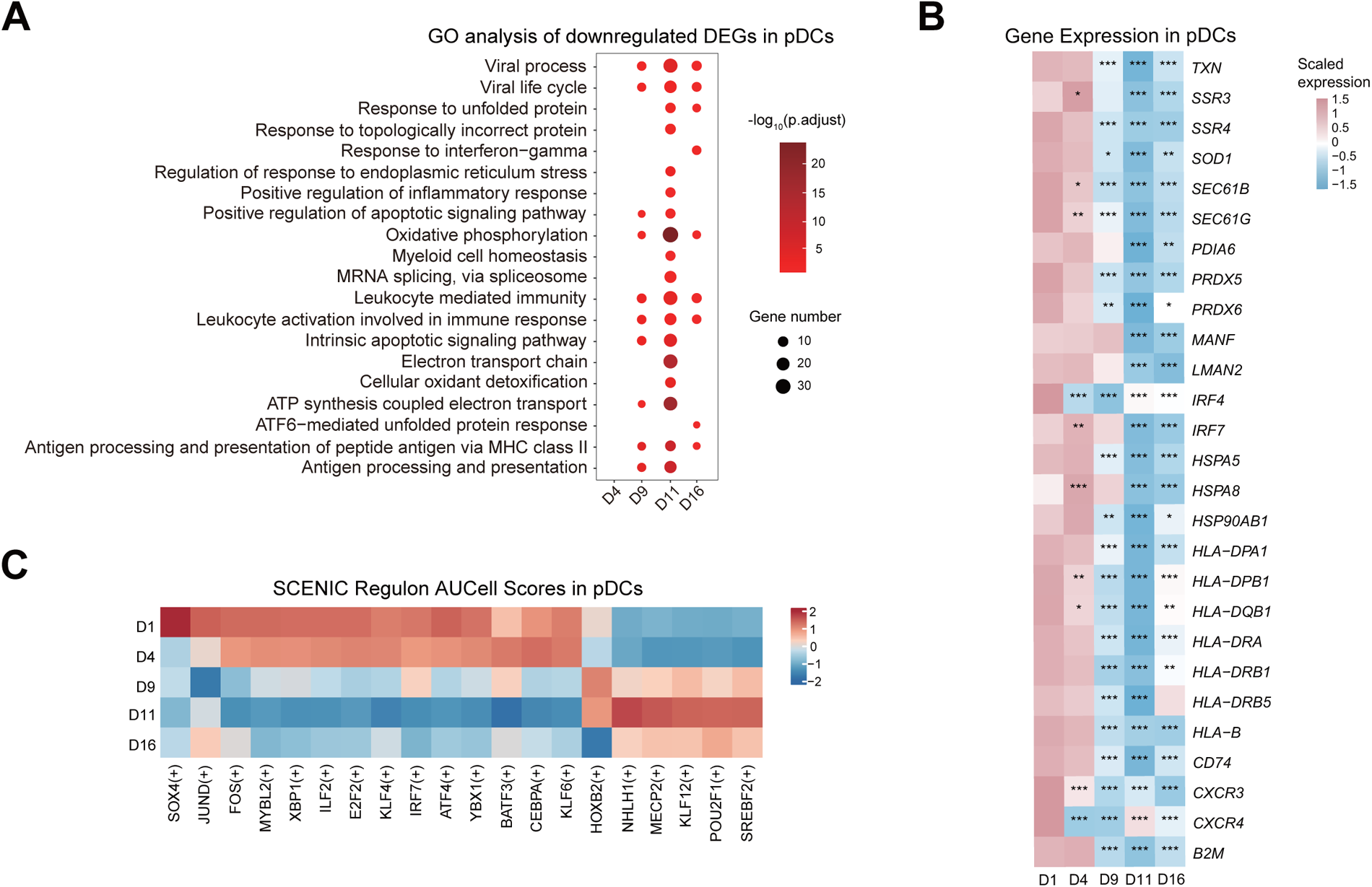
Transcriptional changes in pDCs during mountaineering. (A) The twenty enriched biological processes identified by GO analysis of downregulated DEGs in pDCs. The dot color indicates the statistical significance of the enrichment, and the dot size represents the ratio of genes annotated to each term. (B) Heatmap illustrating scaled expression values of selected DEGs at various time points in pDCs. The two-sided *p* values from the Wilcoxon rank-sum test are shown, \**p* < 0.05, \*\**p* < 0.01, \*\*\**p* < 0.001. (C) Heatmap showing the regulon scenic scores for pDCs across different time points, calculated using the SCENIC algorithm.

**Supplementary Figure 7.**
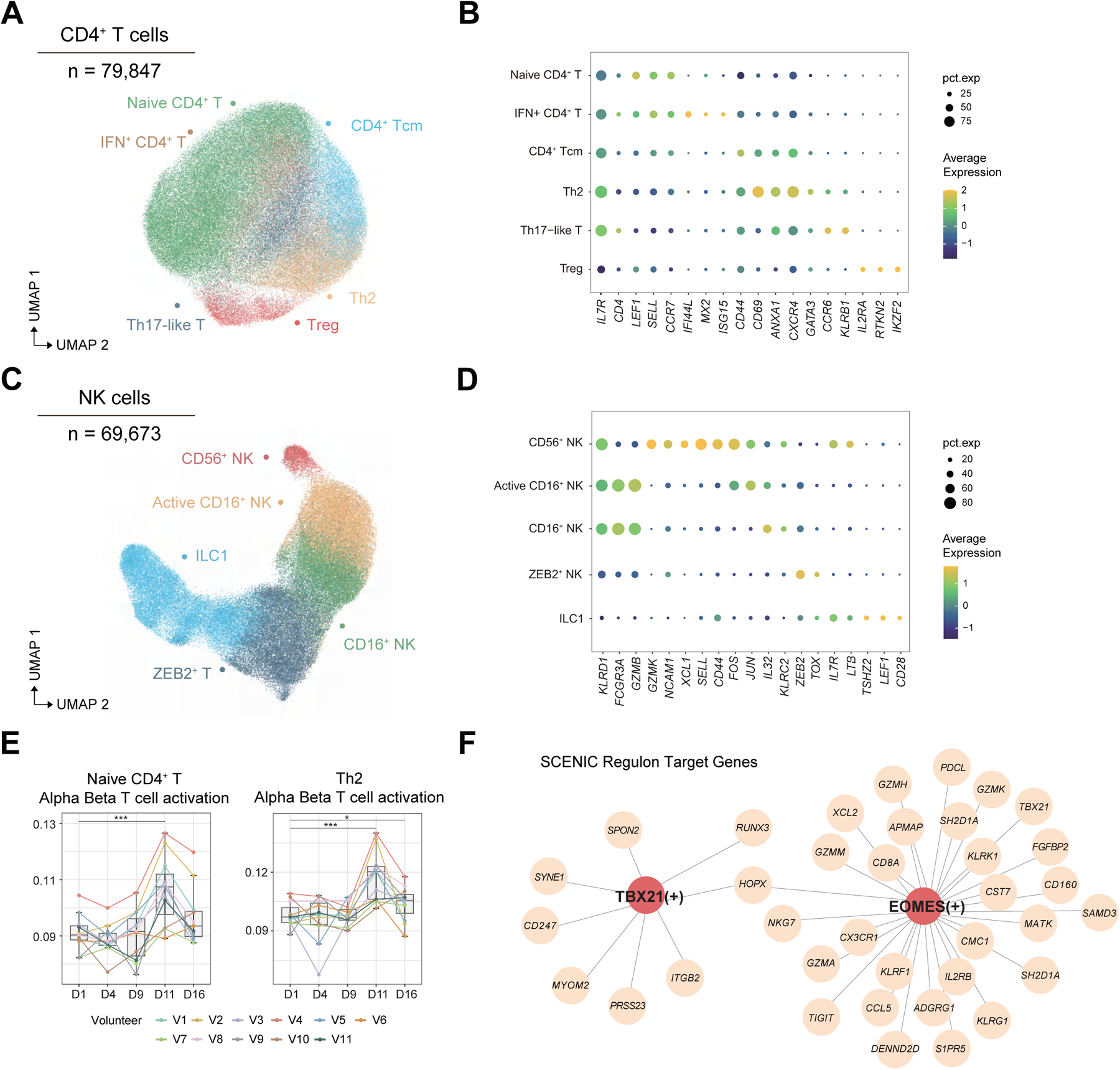
Changes in transcription in CD4^+^ T, CD8^+^ T and NK cell subsets during mountaineering. (A) UMAP plot showing six subsets of CD4^+^ T cells. (B) Dot plot depicting the percentages and average expression levels of canonical genes in six subsets of CD4^+^ T cells. (C) UMAP plot showing five NK cell subsets. (D) Dot plot depicting the percentages and average expression levels of canonical genes in five NK cell subsets. (E) Box plots showing the changes in the alpha/beta T cell activation pathway over time in naive CD4^+^ T and Th2 cells; sample points from the same mountain climber are connected with the same color. The two-sided *p* values from the paired Wilcoxon rank-sum test are shown, \**p* < 0.05, \*\**p* < 0.01, \*\*\**p* < 0.001. (F) The regulatory networks of *TBX21* and *EOMES* and their target genes in CD8^+^ T cells.

**Supplementary Figure 8.**
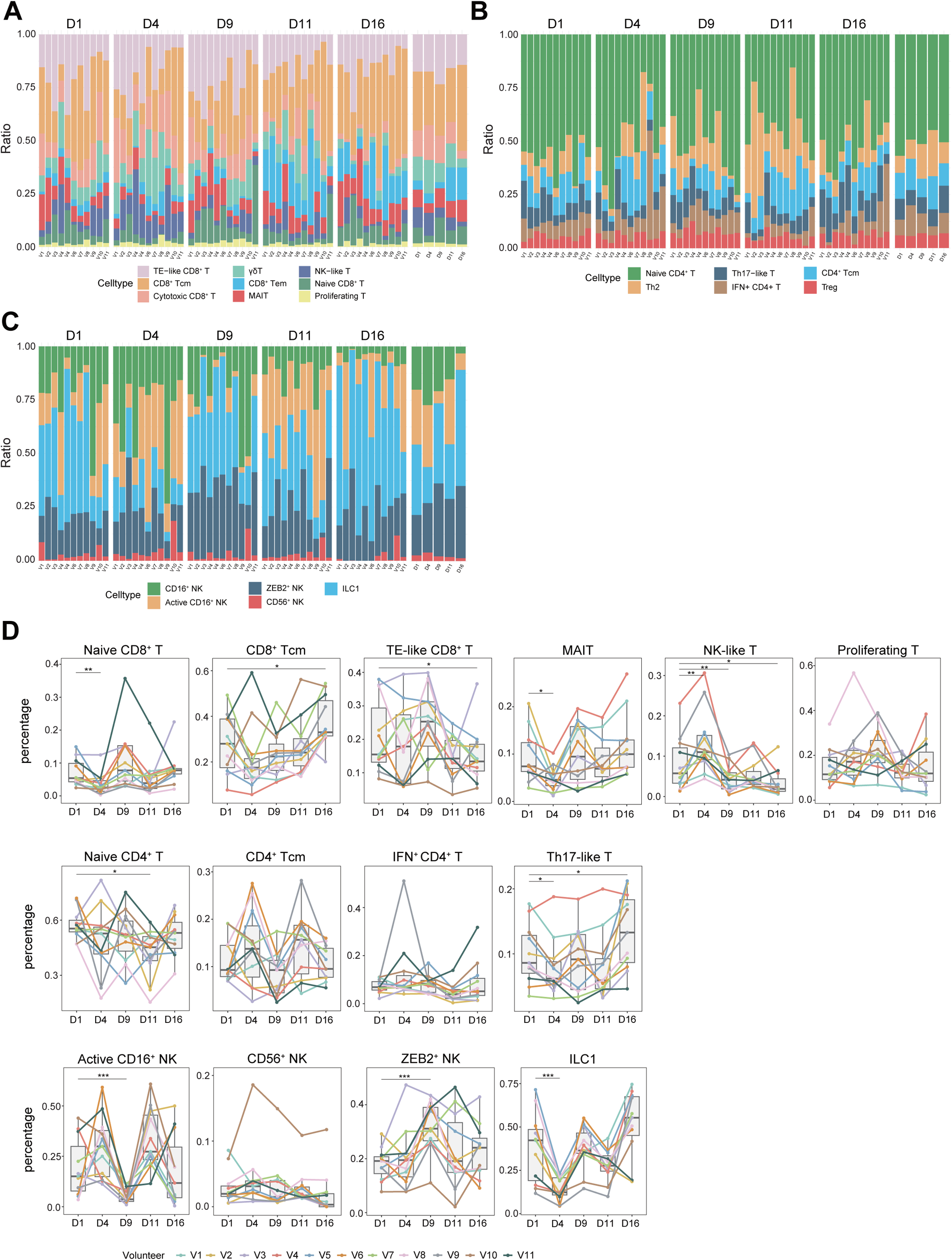
Dynamic changes in the proportions of CD4^+^ T, CD8+ T and NK cell subsets in mountain climbers. (A) Stacked column charts showing the proportions of CD4^+^ T cell subsets from each volunteer and different sampling time points. (B) Stacked column charts showing the proportions of CD8^+^ T cell subsets from each volunteer and at different sampling time points. (C) Stacked column charts showing the proportions of NK cell subsets from each volunteer and different sampling time points. (D) Boxplots illustrating the relative proportions of identified subsets of CD4^+^ T, CD8^+^ T and NK cells during mountaineering (day 1, day 4, day 7, day 11 and day 16, n = 11); sample points from the same mountain climber are connected in the same color. The two-sided *p* values from the paired Wilcoxon rank-sum test are shown, \**p* < 0.05, \*\**p* < 0.01, \*\*\**p* < 0.001.

**Supplementary Figure 9.**
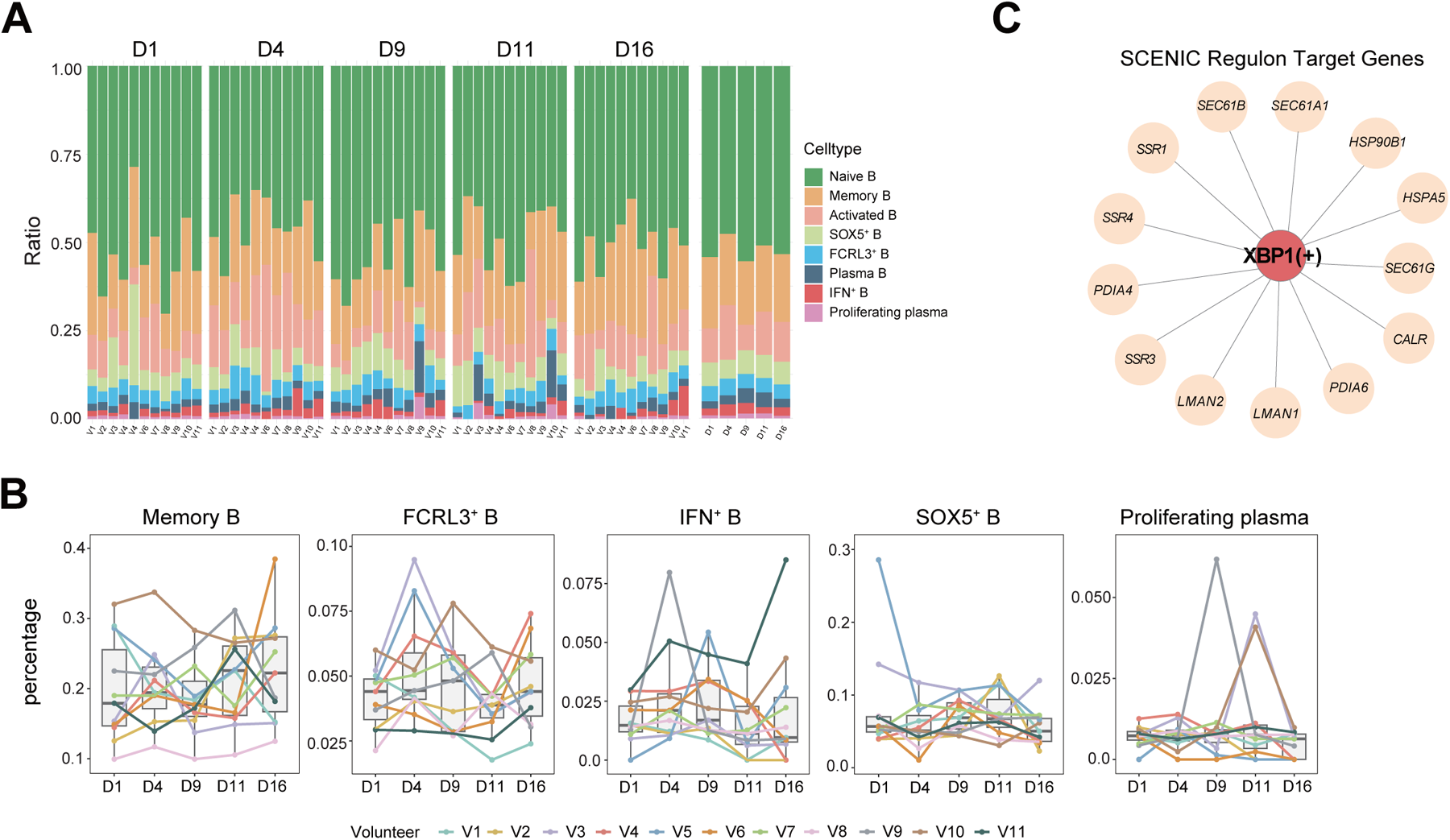
Dynamic changes in the compositions of B cell subsets in mountain climbers. (A) Stacked column charts showing the proportions of B cell subsets from each volunteer and different sampling time points. (B) Boxplots showing the relative proportions of B cell subsets during mountaineering (day 1, day 4, day 7, day 11 and day 16, n = 11); sample points from the same mountain climber are connected with the same color. The two-sided *p* values from the paired Wilcoxon rank-sum test are shown, \**p* < 0.05, \*\**p* < 0.01, \*\*\**p* < 0.001. (C) Regulatory networks of *XBP1* and its target genes in plasma B cells.

**Supplementary Figure 10.**
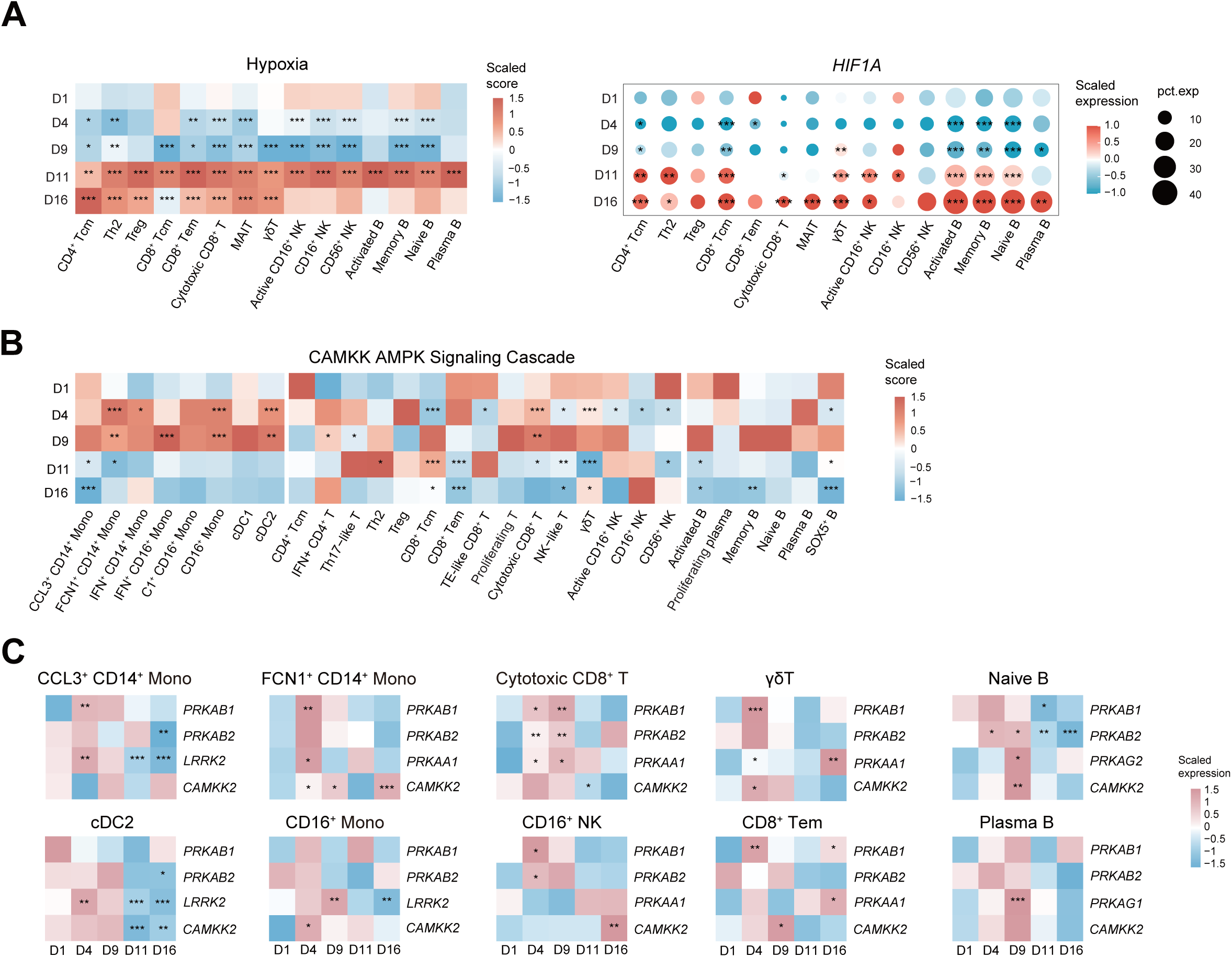
Metabolic pathway analysis of circulating immune cells in mountain climbers. (A) Heatmap showing the scaled expression of genes in the hypoxia pathway and dot plots illustrating the scaled expression and percentages of *HIF1A* in the T, NK and B cell subgroups across different time points. The two-sided *p* values from the Wilcoxon rank-sum test are shown, \**p* < 0.05, \*\**p* < 0.01, \*\*\**p* < 0.001. (B) Heatmap showing the scaled expression of genes in AMPK-related pathways (‘CAMKK AMPK Signaling Cascade’) across different time points. The two-sided *p* values from the Wilcoxon rank-sum test are shown, \**p* < 0.05, \*\**p* < 0.01, \*\*\**p* < 0.001. (C) Heatmap showing the expression levels of AMPK-related genes in immune cell subgroups across different time points. The two-sided *p* values from the Wilcoxon rank-sum test are shown, \**p* < 0.05, \*\**p* < 0.01, \*\*\**p* < 0.001.

**Supplementary Figure 11.**
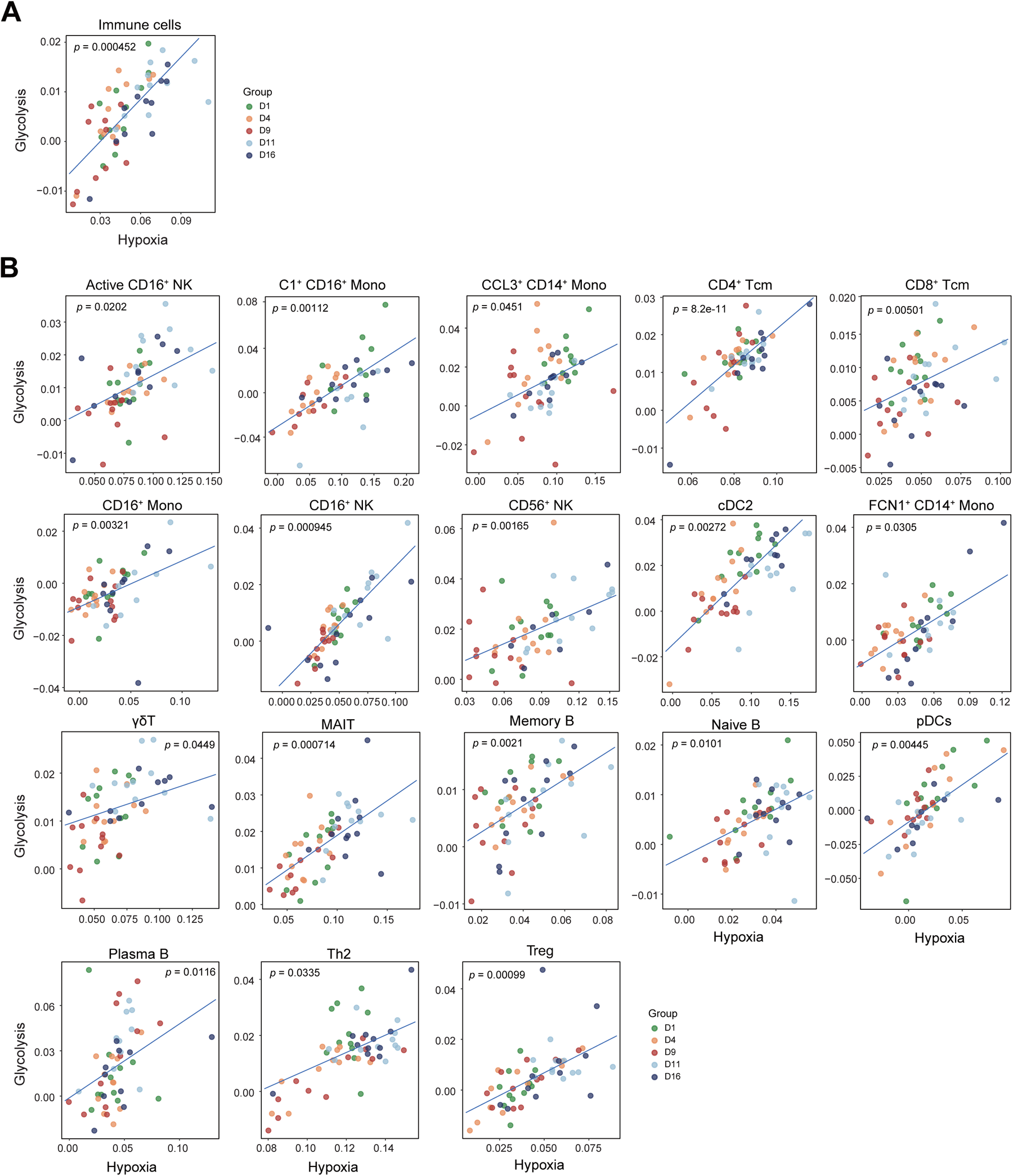
Hypoxia-induced glycolysis pathway during mountaineering. Correlation analysis between hypoxia scores and glycolysis scores in immune cells (A) and each immune subgroup (B) (linear mixed model).

**Supplementary Figure 12.**
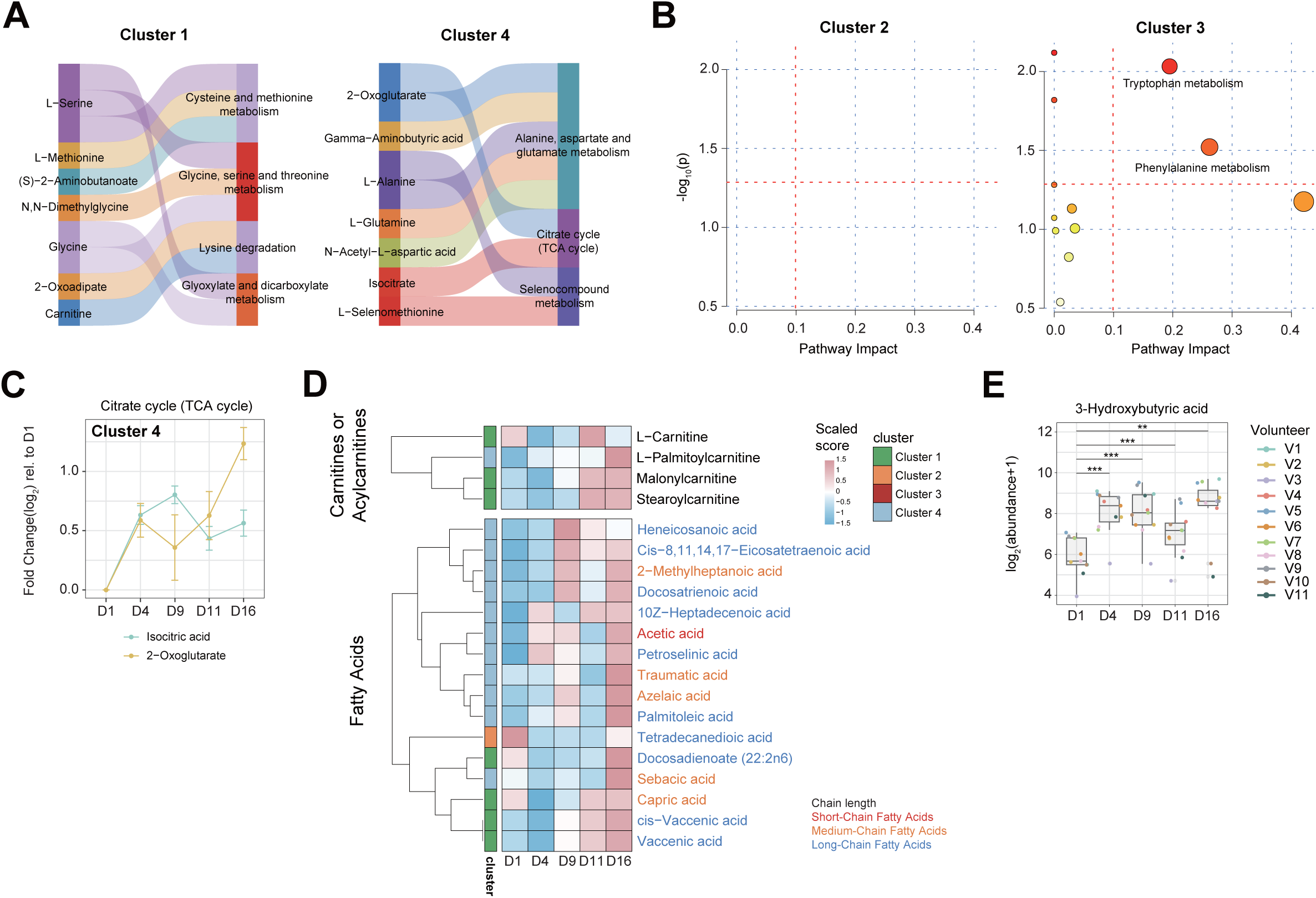
Alterations in plasma metabolites during mountaineering. (A) Sankey diagram illustrating the metabolites contained in metabolic pathways that were enriched in cluster 1 and cluster 4. (B) Pathway enrichment analysis of metabolites from cluster 2 and cluster 3. The red horizontal dashed line indicates the threshold of statistical significance for the y-axis calculated by the hypergeometric test (*p* < 0.05). The red vertical dashed lines indicate pathway impact values ≥ 0.1. (C) Dynamic changes in key metabolites in the citrate cycle (TCA cycle). Dots represent the mean log_2_-fold change relative to D1, and horizontal bars represent the standard error of the mean (SEM). (D) Heatmaps representing changes in the levels of carnitines or acylcarnitines and fatty acids. Clusters are labeled on the left side of the heatmap. Fatty acids with different chain lengths are indicated by different colors. (E) Box plots representing the 3-hydroxybutyric acid levels at different time points; the horizontal line represents the median value. The *p* values from the two-sided Wilcoxon rank-sum test are shown, \**p* < 0.05, \*\**p* < 0.01, \*\*\**p* < 0.001.

## REFERENCES

1. Apollo, M., The true accessibility of mountaineering: The case of the High Himalaya. Journal of Outdoor Recreation and Tourism, 2017. 17: p. 29–43.

2. Luks, A.M. and P.H. Hackett, Medical Conditions and High-Altitude Travel Reply. New England Journal of Medicine, 2022. 386(19): p. 1866–1867.

3. Palmer, B.F., Physiology and Pathophysiology With Ascent to Altitude. American Journal of the Medical Sciences, 2010. 340(1): p. 69–77.

4. Burtscher, J., et al., Immune consequences of exercise in hypoxia: Anarrative review. J Sport Health Sci, 2023.

5. Bosco, G., et al., Body Composition and Endocrine Adaptations to High-Altitude Trekking in the Himalayas. Advancements and Innovations in Health Sciences, 2019. 1211: p. 61–68.

6. Basnyat, B. and J.M. Starling, Infectious Diseases at High Altitude. Microbiol Spectr, 2015. 3(4).

7. Simpson, R.J., et al., Can exercise affect immune function to increase susceptibility to infection? Exercise Immunology Review, 2020. 26: p. 8–22.

8. Nieman, D.C. and L.M. Wentz, The compelling link between physical activity and the body’s defense system. Journal of Sport and Health Science, 2019. 8(3): p. 201–217.

9. Manella, G., et al., The human blood transcriptome exhibits time-of-day-dependent response to hypoxia: Lessons from the highest city in the world. Cell Reports, 2022. 40(7).

10. Rohm, I., et al., Hypobaric hypoxia in 3000 m altitude leads to a significant decrease in circulating plasmacytoid dendritic cells in humans. Clin Hemorheol Microcirc, 2016. 63(3): p. 257–65.

11. Facco, M., et al., Modulation of immune response by the acute and chronic exposure to high altitude. Medicine and Science in Sports and Exercise, 2005. 37(5): p. 768–774.

12. Chen, F., et al., Transcriptome and Network Changes in Climbers at Extreme Altitudes. Plos One, 2012. 7(2).

13. Miller, L.E., et al., Blood Oxidative-Stress Markers During a High-Altitude Trek. International Journal of Sport Nutrition and Exercise Metabolism, 2013. 23(1): p. 65–72.

14. Mrakic-Sposta, S., et al., OxInflammation at High Altitudes: A Proof of Concept from the Himalayas. Antioxidants, 2022. 11(2).

15. Nieman, D.C. and B.D. Pence, Exercise immunology: Future directions. Journal of Sport and Health Science, 2020. 9(5): p. 432–445.

16. Mariggiò, M.A., et al., Peripheral Blood Lymphocytes: A Model for Monitoring Physiological Adaptation to High Altitude. High Altitude Medicine & Biology, 2010. 11(4): p. 333–342.

17. Morabito, C., et al., Responses of peripheral blood mononuclear cells to moderate exercise and hypoxia. Scandinavian Journal of Medicine & Science in Sports, 2016. 26(10): p. 1188–1199.

18. Chi, H.B., Immunometabolism at the intersection of metabolic signaling, cell fate, and systems immunology. Cellular & Molecular Immunology, 2022. 19(3): p. 299–302.

19. O’Brien, K.A., et al., Metabolomic and lipidomic plasma profile changes in human participants ascending to Everest Base Camp. Scientific Reports, 2019. 9.

20. Yu, Y.L., et al., Single-cell sequencing of immune cells after marathon and symptom-limited cardiopulmonary exercise. Iscience, 2023. 26(4).

21. Ehrchen, J.M., et al., The endogenous Toll-like receptor 4 agonist S100A8/S100A9 (calprotectin) as innate amplifier of infection, autoimmunity, and cancer. Journal of Leukocyte Biology, 2009. 86(3): p. 557–566.

22. Swanson, K.V., M. Deng, and J.P.Y. Ting, The NLRP3 inflammasome: molecular activation and regulation to therapeutics. Nature Reviews Immunology, 2019. 19(8): p. 477–489.

23. van de Sande, B., et al., Ascalable SCENIC workflow for single-cell gene regulatory network analysis. Nature Protocols, 2020. 15(7): p. 2247–2276.

24. Pham, K., K. Parikh, and E.C. Heinrich, Hypoxia and Inflammation: Insights From High-Altitude Physiology. Frontiers in Physiology, 2021. 12.

25. Lin, N. and M.C. Simon, Hypoxia-inducible factors: key regulators of myeloid cells during inflammation. Journal of Clinical Investigation, 2016. 126(10): p. 3661–3671.

26. Swiecki, M. and M. Colonna, The multifaceted biology of plasmacytoid dendritic cells. Nature Reviews Immunology, 2015. 15(8): p. 471–485.

27. Iwakoshi, N.N., M. Pypaert, and L.H. Glimcher, The transcription factor XBP-1 is essential for the development and survival of dendritic cells. Journal of Experimental Medicine, 2007. 204(10): p. 2267–2275.

28. Yilmaz, A., et al., Decrease in circulating plasmacytoid dendritic cells during short-term systemic normobaric hypoxia. European Journal of Clinical Investigation, 2016. 46(2): p. 115–122.

29. Wobben, R., et al., Role of hypoxia inducible factor-1α for interferon synthesis in mouse dendritic cells. Biological Chemistry, 2013. 394(4): p. 495–505.

30. Vignali, D.A.A., L.W. Collison, and C.J. Workman, How regulatory T cells work. Nature Reviews Immunology, 2008. 8(7): p. 523–532.

31. Fang, M., et al., CD94 Is Essential for NK Cell-Mediated Resistance to a Lethal Viral Disease. Immunity, 2011. 34(4): p. 579–589.

32. Li, Q., et al., Upregulation of CX3CR1 expression in circulating T cells of systemic lupus erythematosus patients as a reflection of autoimmune status through characterization of cytotoxic capacity. International Immunopharmacology, 2024. 126.

33. Le Bouteiller, P., et al., CD160: A unique activating NK cell receptor. Immunology Letters, 2011. 138(2): p. 93–96.

34. Tu, T.C., et al., CD160 is essential for NK-mediated IFN-γ production. Journal of Experimental Medicine, 2015. 212(3): p. 415–429.

35. Li, M.O. and R.A. Flavell, TGF-β:: A master of all T cell trades. Cell, 2008. 134(3): p. 392–404.

36. Knox, J.J., et al., Characterization of T-bet and eomes in Peripheral Human immune Cells (vol 5, 217, 2014). Frontiers in Immunology, 2016. 7.

37. Krzyzak, L., et al., CD83 Modulates B Cell Activation and Germinal Center Responses. Journal of Immunology, 2016. 196(9): p. 3581–3594.

38. Ricci, D., T. Gidalevitz, and Y. Argon, The special unfolded protein response in plasma cells*. Immunological Reviews, 2021. 303(1): p. 35–51.

39. Chohan, I.S., et al., Immune response in human subjects at high altitude. Int J Biometeorol, 1975. 19(3): p. 137–43.

40. Reimold, A.M., et al., Plasma cell differentiation requires the transcription factor XBP-1. Nature, 2001. 412(6844): p. 300–307.

41. Romero-Ramirez, L., et al., XBP1 is essential for survival under hypoxic conditions and is required for tumor growth. Cancer Research, 2004. 64(17): p. 5943–5947.

42. Lee, P., N.S. Chandel, and M.C. Simon, Cellular adaptation to hypoxia through hypoxia inducible factors and beyond. Nature Reviews Molecular Cell Biology, 2020. 21(5): p. 268–283.

43. Tanner, L.B., et al., Four Key Steps Control Glycolytic Flux in Mammalian Cells. Cell Systems, 2018. 7(1): p. 49-+.

44. Krzeslak, A., et al., Expression of GLUT1 and GLUT3 Glucose Transporters in Endometrial and Breast Cancers. Pathology & Oncology Research, 2012. 18(3): p. 721–728.

45. Herzig, S. and R.J. Shaw, AMPK: guardian of metabolism and mitochondrial homeostasis. Nature Reviews Molecular Cell Biology, 2018. 19(2): p. 121–135.

46. Rolf, J., et al., AMPKα1: A glucose sensor that controls CD8 T-cell memory. European Journal of Immunology, 2013. 43(4): p. 889–896.

47. Wang, J., et al., The regulation effect of AMPK in immune related diseases. Science China-Life Sciences, 2018. 61(5): p. 523–533.

48. Cui, Y.X., et al., The role of AMPK in macrophage metabolism, function and polarisation. Journal of Translational Medicine, 2023. 21(1).

49. Blagih, J., et al., The Energy Sensor AMPK Regulates T Cell Metabolic Adaptation and Effector Responses In Vivo. Immunity, 2015. 42(1): p. 41–54.

50. Halliwell, B., Understanding mechanisms of antioxidant action in health and disease. Nature Reviews Molecular Cell Biology, 2024. 25(1): p. 13–33.

51. Flynn, J.M. and S. Melov, SOD2 in mitochondrial dysfunction and neurodegeneration. Free Radical Biology and Medicine, 2013. 62: p. 4–12.

52. Kaspar, J.W., S.K. Niture, and A.K. Jaiswal, Nrf2:INrf2 (Keap1) signaling in oxidative stress. Free Radical Biology and Medicine, 2009. 47(9): p. 1304–1309.

53. Lee, S., S.M. Kim, and R.T. Lee, Thioredoxin and Thioredoxin Target Proteins: From Molecular Mechanisms to Functional Significance. Antioxidants & Redox Signaling, 2013. 18(10): p. 1165–1207.

54. Zito, E., PRDX4, an Endoplasmic Reticulum-Localized Peroxiredoxin at the Crossroads Between Enzymatic Oxidative Protein Folding and Nonenzymatic Protein Oxidation. Antioxidants & Redox Signaling, 2013. 18(13): p. 1666–1674.

55. Adeva-Andany, M.M., et al., Metabolic effects of glucagon in humans. J Clin Transl Endocrinol, 2019. 15: p. 45–53.

56. Tipton, K.D., D.L. Hamilton, and I.J. Gallagher, Assessing the Role of Muscle Protein Breakdown in Response to Nutrition and Exercise in Humans. Sports Medicine, 2018. 48: p. S53–S64.

57. Lushchak, V.I., Glutathione homeostasis and functions: potential targets for medical interventions. J Amino Acids, 2012. 2012: p. 736837.

58. Yang, L., S. Venneti, and D. Nagrath, Glutaminolysis: A Hallmark of Cancer Metabolism. Annu Rev Biomed Eng, 2017. 19: p. 163–194.

59. Liu, D., et al., Moderate altitude exposure impacts host fasting blood glucose and serum metabolome by regulation of the intestinal flora. Science of the Total Environment, 2023. 905.

60. Cruzat, V., et al., Glutamine: Metabolism and Immune Function, Supplementation and Clinical Translation. Nutrients, 2018. 10(11).

61. Jin, Z., S.K. Mendu, and B. Birnir, GABA is an effective immunomodulatory molecule. Amino Acids, 2013. 45(1): p. 87–94.

62. Schrauzer, G.N., The nutritional significance, metabolism and toxicology of selenomethionine. Adv Food Nutr Res, 2003. 47: p. 73–112.

63. Ayala, A., M.F. Munoz, and S. Arguelles, Lipid peroxidation: production, metabolism, and signaling mechanisms of malondialdehyde and 4-hydroxy-2-nonenal. Oxid Med Cell Longev, 2014. 2014: p. 360438.

64. Makrecka-Kuka, M., et al., Plasma acylcarnitine concentrations reflect the acylcarnitine profile in cardiac tissues. Sci Rep, 2017. 7(1): p. 17528.

65. Nelson, A.B., et al., Metabolic Messengers: ketone bodies. Nat Metab, 2023. 5(12): p. 2062–2074.

66. Tsekouras, Y.E., et al., A single bout of whole-body resistance exercise augments basal VLDL-triacylglycerol removal from plasma in healthy untrained men. Clin Sci (Lond), 2009. 116(2): p. 147–56.

67. Liu, P., et al., The mechanisms of lysophosphatidylcholine in the development of diseases. Life Sci, 2020. 247: p. 117443.

68. van der Veen, J.N., et al., The critical role of phosphatidylcholine and phosphatidylethanolamine metabolism in health and disease. Biochimica Et Biophysica Acta-Biomembranes, 2017. 1859(9): p. 1558–1572.

69. Nieman, D.C., et al., Serum Metabolic Signatures Induced By a Three-Day Intensified Exercise Period Persist After 14 h of Recovery in Runners. Journal of Proteome Research, 2013. 12(10): p. 4577–4584.

70. Iqbal, J., et al., Sphingolipids and Lipoproteins in Health and Metabolic Disorders. Trends in Endocrinology and Metabolism, 2017. 28(7): p. 506–518.

71. Da Rosa, P.C., et al., The physical exercise-induced oxidative inflammatory response in peripheral blood mononuclear cells: Signaling cellular energetic stress situations. Life Sciences, 2023. 321.

72. Laddu, R., et al., Physical activity for immunity protection: Inoculating populations with healthy living medicine in preparation for the next pandemic. Progress in Cardiovascular Diseases, 2021. 64: p. 102–104.

73. Pham, K., et al., Inflammatory gene expression during acute high-altitude exposure. Journal of Physiology-London, 2022. 600(18): p. 4169–4186.

74. Spaulding, H.R. and Z. Yan, AMPK and the Adaptation to Exercise. Annual Review of Physiology, 2022. 84: p. 209–227.

75. Kammerer, T., et al., Hypoxic-Inflammatory Responses under Acute Hypoxia: In Vitro Experiments and Prospective Observational Expedition Trial. International Journal of Molecular Sciences, 2020. 21(3).

76. McNicholas, W.T., Chronic Obstructive Pulmonary Disease and Obstructive Sleep Apnea Overlaps in Pathophysiology, Systemic Inflammation, and Cardiovascular Disease. American Journal of Respiratory and Critical Care Medicine, 2009. 180(8): p. 692–700.

77. Rius, J., et al., NF-κB links innate immunity to the hypoxic response through transcriptional regulation of HIF-1α. Nature, 2008. 453(7196): p. 807–U9.

78. Mingyuan, X., et al., Hypoxia-inducible factor-1alpha activates transforming growth factor-beta1/Smad signaling and increases collagen deposition in dermal fibroblasts. Oncotarget, 2018. 9(3): p. 3188–3197.

79. Regis, S., et al., NK Cell Function Regulation by TGF-β-Induced Epigenetic Mechanisms. Frontiers in Immunology, 2020. 11.

80. Murray, A.J. and J.A. Horscroft, Mitochondrial function at extreme high altitude. Journal of Physiology-London, 2016. 594(5): p. 1137–1149.

81. Sinha, S., et al., Different adaptation patterns of antioxidant system in natives and sojourners at high altitude. Respiratory Physiology & Neurobiology, 2009. 167(3): p. 255–260.

82. Dobin, A., et al., STAR: ultrafast universal RNA-seq aligner. Bioinformatics, 2013. 29(1): p. 15–21.

83. Butler, A., et al., Integrating single-cell transcriptomic data across different conditions, technologies, and species. Nature Biotechnology, 2018. 36(5): p. 411-+.

84. McGinnis, C.S., L.M. Murrow, and Z.J. Gartner, DoubletFinder: Doublet Detection in Single-Cell RNA Sequencing Data Using Artificial Nearest Neighbors. Cell Systems, 2019. 8(4): p. 329-+.

85. Yu, G.C., et al., clusterProfiler: an R Package for Comparing Biological Themes Among Gene Clusters. Omics-a Journal of Integrative Biology, 2012. 16(5): p. 284–287.

86. Liberzon, A., The Molecular Signatures Database (MSigDB) hallmark gene set collection. Cell systems, 2015.

87. Tyanova, S., et al., The Perseus computational platform for comprehensive analysis of (prote)omics data. Nature Methods, 2016. 13(9): p. 731–740.

88. Pang, Z.Q., et al., Using MetaboAnalyst 5.0 for LC-HRMS spectra processing, multi-omics integration and covariate adjustment of global metabolomics data. Nature Protocols, 2022. 17(8): p. 1735–1761.

89. Mfuzz: a software package for soft clustering of microarray data. 2007.

90. Lee, J.W., et al., Integrated analysis of plasma and single immune cells uncovers metabolic changes in individuals with COVID-19. Nature Biotechnology, 2022. 40(1): p. 110-+.

91. Bolker, B.M., et al., Generalized linear mixed models: a practical guide for ecology and evolution. Trends in Ecology & Evolution, 2009. 24(3): p. 127–135.

